# Towards optimizing diversifying base editors for high-throughput studies of single- nucleotide variants

**DOI:** 10.1101/2024.11.18.621003

**Authors:** Carley I Schwartz, Nathan S Abell, Amy Li, Aradhana, Josh Tycko, Alisa Truong, Stephen B Montgomery, Gaelen T Hess

**Author notes:** These authors contributed equally to this work (Carley I Schwartz, Nathan S Abell).

## Abstract

Determining the phenotypic effects of single nucleotide variants is critical for understanding the genome and interpreting clinical sequencing results. Base editors, including diversifying base editors that create C>N mutations, are potent tools for installing point mutations in mammalian genomes and studying their effect on cellular function. Numerous base editor options are available for such studies, but little information exists on how the composition of the editor (deaminase, recruitment method, and fusion architecture) affects editing. To address this knowledge gap, the effect of various design features, such as deaminase recruitment and delivery method (electroporation or lentiviral transduction), on editing was assessed across ∼200 synthetic target sites. The direct fusion of a hyperactive variant of activation-induced cytidine deaminase to the N-terminus of dCas9 (DivA-BE) produced the highest editing efficiency, ∼4-fold better than the previous CRISPR-X method. Additionally, DivA-BE mutagenized the DNA strand that anneals to the targeting sgRNA to create G>N mutations, which were absent when the deaminase was fused to the C-terminus of dCas9. The DivA-BE editors efficiently diversified their target sites, an ideal characteristic for discovering functional variants. These and other findings provide a comprehensive analysis of how design features influence the activity of several popular base editors.

## Introduction

Advances in sequencing technologies have enabled the detection of single nucleotide variants (SNVs) in patients. However, the benefit of this wealth of information will only be realized when the effect of any given SNV on gene or cellular function is known. Most identified SNVs, even those in disease-associated genes, are classified as variants of unknown significance (VUS) (1), which means they cannot be acted on clinically. With the average human genome containing 4- 5 million SNVs, there is an urgent need for tools and methods that will enable the accurate and high-throughput assessment of the impact of SNVs.

Base editors are potent genome editors that can introduce SNVs in genomic DNA, allowing the effect of individual base substitutions in cell or animal models to be studied. As these editors do not generate toxic DNA double-strand breaks, they reduce the introduction of insertions/deletions (indels) that can result in unwanted gene knock-outs. The initial Cas9-mediated base editors generated C>T substitutions by recruiting a cytidine deaminase and bacteriophage uracil glycosylase inhibitor (UGI) to a genomic site via a nickase Cas9 (nCas9) (2–4). Since this initial demonstration of the approach, there have been a plethora of additional editors that use engineered deaminases, including ones that modify adenines (5). These editors have been used to assess the effect of SNVs on gene function in a multitude of contexts (6).

In addition to these programmable C>T base editors, diversifying cytidine base editors (DCBEs) that produce C>N edits are powerful tools for annotating single nucleotide variants (SNVs) in high-throughput assays. Beyond these clinical applications, DCBEs have enabled the discovery of mutations that promote drug resistance and have assisted the design of novel proteins *in situ* (7–9). These diversifying base editors hold an advantage over C>T editors as they produce more substitutions that can be interrogated for phenotypic effects. For diversifying and programmable base editors, characterization of their properties (efficiency, targeting window, the spectrum of substitutions, etc.) is necessary for selecting the optimal editor for studying SNV function. Furthermore, there can be a trade-off between these editing properties. For example, increases in editing efficiency can expand the base editing window, which may not be desirable for making a specific base substitution.

Given the number of base editor options available for these SNV studies, it is critical to define the properties of base editors to determine the editor most suited for the intended application. Understanding how the base editor configuration (e.g., method of recruiting or fusing the deaminase to the Cas9 protein) affects editing properties would facilitate the creation of editors better suited for their specific studies. Defining the properties of editors would benefit from the parallel assessment of the editor’s activity across many target sites, as underlying sequence variation can influence editing outcomes. While such studies have profiled programmable editors (10, 11), to date, studies of DCBEs have focused on editing for a limited number of target sequences (7, 8, 12).

Here, a comprehensive approach was established and used to assess the impact of several design features on the editing outcomes of DCBEs for ∼200 sgRNA-target site pairs simultaneously. We determined the editing efficiency, targeting window, and base substitution spectrum for base editors expressed transiently or constitutively. The recruitment method for a hyperactive variant of the activation-induced cytidine deaminase AID affected the editing efficiency and targeting window. These studies established a new diversifying base editor, DivA- BE, consisting of a hyperactive AID variant fused to the N-terminus of catalytically inactive Cas9 (dCas9) with improved editing efficiency and a reduced targeting window compared to the previously established CRISPR-X editor (7). Additionally, G>N mutations were observed for DivA- BE but were depleted for editors with the AID variant fused to the C-terminus of dCas9 and editors using the rat APOBEC1 (rAPOBEC1) deaminase. Our analyses revealed that fusing the hyperactive AID variant to nCas9 (nDivA-BE) produced the most diversification at the target site but also exhibited the highest frequency of indels, which may be a drawback in assays testing the phenotypic effect of SNVs. These and other data described in this report should serve as an important resource for maximizing the effectiveness of future high throughput editing experiments.

## Materials and Methods

### Design of sgRNA-target site library

To design the sgRNA library, we selected 2000 20-bp sgRNA sequences from non-targeting controls generated for a human genome-wide CRISPR knockout library (13). The target site was 104 bp long, centered on an NGG PAM compatible with editing by *Streptococcus pyogenes* Cas9 (Cas9). The N in the NGG and the surrounding sequence beyond the sgRNA were randomly generated with a 52% GC bias to represent the content of the human genome. For half of the sites, the sequence was reverse-complemented. sgRNAs and target sites containing restriction enzyme sites (BsmBI/Esp3I, BsaI, and BlpI) required for cloning were removed, and 200 sgRNA-target site pairs were selected randomly. For ten sgRNA-target site pairs, the sgRNA sequence in the target site was scrambled to produce negative control sgRNA-target site pairs. A linker sequence (5’-GTTTGGAGACGGGATACCGTCTCTGATC-3’) containing two BsmBI sites was placed after the sgRNA protospacer sequence and before the target site. This linker sequence was used to insert the sgRNA stem-loop during cloning. The oligonucleotides were extended on both ends to add sequences to enable PCR amplification of the oligonucleotides and add restriction sites compatible with cloning at the 5’ (BsaI) and 3’ (BlpI) sites, respectively. A complete list of the designed oligonucleotides are listed in **Supplementary Table 1**. These designed oligonucleotides were synthesized by Agilent.

### Construction of paired sgRNA-target site library

The sequences of primers used to generate the sgRNA-target site library are listed in **Supplementary Table 2.** The sgRNA-target site oligonucleotides were cloned into pGH643. This vector contained a human U6 promoter expressing a sgRNA upstream of an Ef1ɑ promoter driving the expression of puromycin resistance and mCherry used for downstream selection and monitoring infection percentage. A 100 μL restriction digest of pGH643 was incubated overnight at 37°C (10 μg pGH643, 4.5 μL Esp3I (NEB R0734L), 4.5 μL BlpI (NEB R0585L), and 10 μL of CutSmart Buffer (NEB)). The digest reaction was run on a 1% TAE gel, and the band was excised and purified with the QiaQuick Gel Extraction Kit (Qiagen 28706). The oligonucleotide library insert was produced using 4 × 100 μL PCR reactions. Each reaction contained 1.5 ng (synthesized oligonucleotide template), 1 μL of Herculase II polymerase (Agilent 600679), 2 μL 10 mM dNTPs (Roche 11969064001), 2 μL 100 μM forward primer, 2 μL 100 μM reverse primer, 2 μL DMSO, and 20 μL of 5× Herculase buffer. The thermocycling conditions were 98°C for 3 min, 11 cycles of 98°C 30 sec, 55°C for 30 sec, 72°C for 30 sec, followed by a final extension of 72°C for 3 min. The reactions were pooled and 5 μL was analyzed on a 2% TBE agarose gel to confirm the correct amplicon size. The remaining PCR reaction was purified using the QiaQuick PCR Purification Kit (Qiagen 28104) and eluted in EB buffer. The purified PCR was digested overnight at 37°C in a 120 μL reaction (5 μL BsaI-HF (NEB R3733L), 5 μL BlpI, and 12 μL of CutSmart Buffer). The digested insert was analyzed on a 3% TBE agarose gel and bands were extracted and purified. A 100 μL ligation reaction was set up as follows: 2 μg digested pGH643, 360 ng digested insert (∼12:1 ratio), 5 μL of T4 Ligase (NEB M0202L), and 10 μL T4 Ligase Buffer (NEB B0202S). The reaction was incubated for 16 hr at 16°C followed by 70°C for 15 min. The ligation reaction was purified and concentrated to >100 ng/μL using MinElute PCR Purification Kit (Qiagen 28006). 200 ng of purified ligation was electroporated into 60 μL of electrocompetent Lucigen Endura cells (Biosearch Technologies 60242-1) according to manufacturer instructions. Cells were rescued in 4 mL of Lucigen recovery media and grown for 1 hr at 37°C. The library was plated on 8 × 1 sq ft. LB agar (Fisher DF0445076) plates with carbenicillin (Fisher 50213248, 100 μg/mL). Two smaller plates were used to titer the number of transformed colonies and guarantee the library had > 100× coverage per sgRNA-target site pair. The 1 sq ft. LB agar plates were scraped to collect colonies, and DNA was extracted using the HiSpeed Plasmid Maxi Kit (Qiagen 12663).

A second round of cloning was performed to introduce the stem-loop between the sgRNA protospacer and the target site. The no stem-loop plasmid library was digested for 8 hr at 55°C (10 μg plasmid library, 9 μL BsmBI (NEB R0739L), and 10 μL of Buffer 3.1 (NEB B6003S). The resulting digest was analyzed on a 1% TAE agarose gel and purified. The stem-loop insert was prepared by 4×100 PCR reactions consisting of 1 ng of pGH643 (stem-loop template), 1 μL of Herculase II polymerase, 2 μL 10 mM dNTPs, 2 μL 100 μM oGH915, 2 μL 100 μM oGH1001, 2 μL DMSO, and 20 μL of 5× Herculase buffer. oGH1001 included 7 degenerate nucleotides that serve as a unique molecular identifier (UMI). This UMI was inserted between the end of the stem-loop and the start of the target site. The thermocycling conditions were 98°C for 3 min, 15 cycles of 98°C 30 sec, 55°C for 30 sec, 72°C for 30 sec, followed by a final extension of 72°C for 3 min. The fragment was analyzed on a 2% TBE agarose gel, followed by gel extraction and purification. The purified PCR product was digested for 8 hr at 55°C in a 120 μL reaction (9 μL BsmBI and 12 μL Buffer 3.1), followed by PCR purification. A 100 μL ligation reaction consisting of 2 μg of digested library, 280 ng digested stem-loop (∼10:1 ratio), 5 μL T4 Ligase, and 10 μL T4 Ligase Buffer was incubated at 16°C for 16 hr. The reaction was purified and concentrated using the MinElute PCR Purification Kit to a concentration > 100 ng/μL. 200 ng of the cleaned ligation was electroporated into Lucigen Endura cells and plated as described for the initial plasmid library. DNA was extracted from bacteria using the Qiagen HiSpeed Plasmid Kit.

### Construction of base editor expression vectors

A complete list of plasmids and their sequences are listed in **Supplementary Table 2**. Expression constructs for MS2-AID variant fusions (pGH335, pGH183) and dCas9 (pGH125) were previously described (7). pGH335 and pGH183 contained an Ef1ɑ promoter driving the expression of the MS2 coat protein fused to an AID variant. These vectors were upstream of a T2A-hygromycin resistance (HygroR) cassette. Both vectors were digested with BsrGI-HF (NEB R3575L) and EcoRI-HF (NEB R3101L) to remove the HygroR. A PCR-amplified T2A-GFP cassette was inserted in place of HygroR by Gibson Assembly. pGH125 contained an Ef1ɑ promoter driving the expression of dCas9 followed by a T2A-blasticidin resistance (BlastR) gene. A fluorescent dCas9-expression vector (pGH483) was generated by digesting pGH125 with BamHI-HF (NEB R3136L) and EcoRI-HF to remove T2A-BlastR. A T2A-tagBFP cassette was amplified by PCR and inserted into the digested pGH125 via Gibson assembly. pGH125 or pGH483 were digested with BsiWI-HF (NEB R3553L) and AfeI (NEB R0652S) to remove dCas9, and genome editors were inserted into the vector via Gibson Assembly. pGH896, which expressed DivA-BE, was digested with BamHI-HF and EcoRI-HF to remove the T2A-tagBFP, and T2A-GFP was inserted in its place to generate pGH941.

### Cell Culture

K562 cells, a chronic myeloid leukemia model, were cultured in RPMI 1640 media (Gibco 11875119) supplemented with 10% FBS (Life Technologies 10437028), 2 mM L-glutamine (VWR 16777-162), and 1% penicillin/streptomycin (VWR 16777-164). HEK293T cells were grown in DMEM (Gibco 11995065) with 10% FBS, 2 mM L-glutamine, and 1% penicillin/streptomycin. All cell lines were maintained in a humidified incubator (37°C, 5% CO_2_) and checked regularly for mycoplasma contamination.

### Generating paired sgRNA-target site library in cell lines

Lentivirus for the sgRNA-target site library was produced as follows. 5×10^5^ HEK293T cells were plated in all wells of a 6-well plate in 2 mL of DMEM and incubated overnight. Each well was transfected with 750 ng plasmid library, 750 ng of third-generation packaging mix (equimolar mix of pMD2.G (Addgene #12259), psPAX2 (Addgene #12260), and pMDLg/pRRE (Addgene #12251)), and 10 μL polyethylenimine (PEI, Polysciences Inc. 239661, 1 mg/mL). 24 hr after transfection, 3 mL of DMEM is added to each well. Lentivirus was harvested 72 hr after transfection and filtered through a 0.45 μm syringe filter to remove debris. The virus from each well was pooled, resulting in ∼30 mL of virus. 10 mL of virus at three dilutions (1:2, 1:5, and 1:10) were used to infect 10^6^ wild-type K562 cells for 2 hr at 1000 *g* at 33°C in duplicate. Polybrene (Sigma-Aldrich H92685G, 8 μg/mL) was added to the viral media. After centrifugation, cells were resuspended in regular culture media and grown for 3 days. The transduction efficiency was evaluated for each dilution via flow cytometry of mCherry percentage. We chose the 1:5 dilution of lentivirus, which had a transduction efficiency of ∼10%, which suggests that most cells contained only a single sgRNA-target site integrated. The infected cells were selected with puromycin (Sigma-Aldrich P8833, 1 μg/mL) for 5 days until greater than 90% of the cells were mCherry positive (mCherry+). After puromycin selection, the cells were bottlenecked to reduce the complexity of the sgRNA-target site library. 10^5^ mCherry+ K562 cells were plated and expanded to grow until the introduction of base editors.

### Induction of base editing in library cells

For electroporated samples, 5 × 10^6^ bottlenecked K562 cells were centrifuged at 500 *g* for 5 min. Cells were resuspended in 100 μL of electroporation buffer. The electroporation buffer consisted of 20 μL of Buffer I (0.2 g ATP disodium salt (Sigma-Aldrich A6419-10G) and 0.12 g magnesium chloride hexahydrate (Fisher Scientific BP214-500) in 1 mL) and 1 mL of Buffer II (0.06 g potassium dihydrogen phosphate (Sigma-Aldrich P5655-100G), 0.06 g sodium bicarbonate (Sigma-Aldrich S5761-500G), 0.02 g glucose (Sigma-Aldrich G7021-100G) in 50 mL at pH 7.4). 1 μg of each plasmid being electroporated was added to the cell mixture and moved to a 2 mm cuvette. Cells were electroporated on a Lonza Nucleofector 2b using program T-016. Cells were rescued in 3 mL of warm media and allowed to rest for 24 hr. Afterward, cells were expanded and maintained in log growth. 3 days after electroporation, cells were sorted on a BD FACSAria II to collect the desired fluorescent cells (tagBFP positive or tagBFP/GFP positive). The number of cells collected varied between 3-5 × 10^5^ cells. Sorted cells were grown for an additional 6 days, after which cells were collected for sequencing. For the ABE7.10;DivA-BE;Sequential sample, a second round of electroporation was performed on the ABE7.10 edited cells.

For lentivirally transduced samples, lentivirus was produced using the same protocol described for the sgRNA-target site library. 2 mL of lentivirus per plasmid was used to transduce 3 × 10^5^ bottlenecked K562 cells. Polybrene (8 μg/mL) was added to the virus-cell mixture, and the cells were centrifuged at 1000 *g* for 2 hr at 33°C. After centrifugation, the viral media was removed and replaced with fresh RPMI growth media. Cells were grown for 2 days before adding blasticidin (Research Products International, B12200, 10 μg/mL) and/or hygromycin B (Thermo Fisher 10687010, 200 μg/mL). Cells were selected for 6 days before collection for sequencing. Bottlenecked K562 cells not transduced with editors were treated with each antibiotic in parallel to verify that the selection was complete.

### Preparation and sequencing of sgRNA-target site amplicon libraries

Genomic DNA was extracted from 3 × 10^6^ cells using QiaAmp DNA Blood Mini Kits (Qiagen 51104) following the manufacturer’s instructions. The sgRNA-target site-UMI cassette was amplified using PCR with primers listed in **Supplementary Table 2**. These primers added Illumina adapters and sample barcodes. 2 × 100 μL reactions were set up for each sample. Each 100 μL reaction contained 5 μg of genomic DNA, 1 μL of 100 μM Forward primer mix (oGH840.1-6), 1 μL of 100 μM reverse barcoded primer, and 50 μL of NEBNext 2× Master Mix (NEB M0541S). oGH840.1-6 are six oligonucleotides with variable lengths to stagger and phase the sequencing of a common sequence in the target site amplicon. Samples were placed in a thermocycler and heated to 98°C for 3 min, followed by 25 cycles of 98°C for 30 s, 59°C for 30 s, 72°C for 4 min. PCRs were pooled, and 50 μL of this mixture was analyzed on a 2% TBE agarose gel. A band at 430 bp was excised and purified with the QIAquick Gel Extraction Kit. Libraries were quantified with a Qubit dsDNA HS Kit (ThermoFisher Q33231) and pooled evenly except for Parent samples, which were loaded at 2× compared to other samples. PhiX control (Illumina FC-110-3001) was spiked into the mixture of prepared samples at 30% and sequenced on an Illumina NextSeq 550 using a paired-end read of 139 cycles with the standard Read 1 primer, a 6-cycle Index 1 read with a custom primer (oGH778), and a 21-cycle read with a custom Read 2 primer (oGH779).

### Processing of sequencing data files

Sequencing reads were demultiplexed using bcl2fastq. A bowtie (version 1.3.1) (14) (sgRNAs) and bowtie2 (version 2.4.5) (15) (target sites) indices were generated using ‘make_fa_sgRNA_Indices.py’ and ‘make_fa_TargetSite_Indices_b2.py’, respectively. Common sequences were removed, and UMIs were extracted from reads using ‘BEChar_extract_trim.py’. This script uses umi_tools (version 1.1.6) (16) to extract the 7-bp UMI from Read 1. The UMI is appended to each Read_ID by umi_tools. Cutadapt (version 4.9) (17) removed a common primer sequence used to amplify the region from Read 1 after UMI extraction (target site) and the last base from Read 2 (sgRNA).

The fastq files from the Parent samples were aligned, and sgRNA-UMI-triplets to be included in the analysis were generated using ‘Define_whitelist.py’. The sgRNA alignment was done with bowtie, allowing one mismatch in the -v mode. We aligned the target site using bowtie2 in the -- local mode. The resulting sam files were converted to bam files using samtools (version 0.1.19) (18). We compared four read-filtering pipelines (see **Supplementary Text S1**). The final processing pipeline contained three read-filtering steps. In the first step, reads that failed to align to any target site were removed. Second, reads with improperly paired target site and sgRNA were removed. For the last filtering step, umi_tools grouped reads into sgRNA-target site-UMI triplets. We tabulated a list of triplets where >90% of their reads were free of insertions/deletions in the Parent sample for each replicate. This list of acceptable sgRNA-UMI-triplets was used to filter the fastqs for all samples in downstream steps.

### Generating allele table for Transient and Integrated samples

Fastqs generated after UMI extraction and truncation for all Transient and Integrated samples were processed using ‘UMI_Filter.py’. This script filters each fastq file by retaining reads that match a sgRNA-target site-UMI triplet found in our list of acceptable triplets from the Parent sample from that replicate. The subsequent fastqs were processed using CRISPResso2 (version 2.2.11) (19) using the CRISPRessoPooled command with the following settings: -- exclude_bp_from_left 5 --exclude_bp_from_right 0 --min_reads_to_use_region 10 -- base_editor_output --suppress_plots -w 0 --write_detailed_allele_table. We combined all of these target sites and editors into a single allele table using ‘CreateAlleleTables_General.py’ followed by normalization of the tables using ‘NormalizeTables_Cutoff.py’. In the normalization script, alleles with only one read were removed as these alleles are more likely to contain mutations introduced by sequencing error, and the frequency of each allele was recalculated after removing these alleles. The analysis was applied to the Transient and Integrated samples separately, and allele tables were generated for each dataset.

Normalized allele and base substitution frequencies relative to the Parent sample were generated by ‘MakeAllTables_Full.r’ (Transient) and ‘MakeAllTables_Full_Lenti.r’ (Integrated). We calculated the normalized frequency of each allele detected by subtracting the frequency of the same allele in the Parent sample. For alleles not detected in the Parent sample, we set the Parent allele frequency to 0. Any negative frequencies after normalization to the Parent allele frequency were set to 0. If an allele was only detected in one replicate, the normalized frequency in the other replicate was set to 0. The mean editing efficiency for each allele was calculated using the normalized frequency from both replicates. This normalized frequency table was used to create a base substitution frequency table by separating each base substitution observed in a mutated allele. This separation produces a row in the table for each substitution containing its position in the allele and the normalized frequency of the allele from which the substitution is derived.

### Analysis of base editing and indel efficiency

Editing efficiency for each target site was calculated by summing the normalized allele frequency of all modified alleles with a frequency above 0.5% for that target site. Target sites with fewer than 200 reads across both replicates were removed from the analysis. For indel editing efficiency, the normalized allele frequency table was limited to alleles that had n_inserted or n_deleted >0. These two columns represent the number of inserted or deleted bases, as calculated by CRISPResso2. For the calculation of editing efficiency for alleles with G>N or C>N edits, the sequences were reverse complemented for target sites where the sgRNA targeted the bottom strand, converting these target sites to match those with a top strand targeting sgRNA. An additional column was calculated to specify whether the allele contained G>N, C>N, or both types of edits. C>N editing efficiency was determined using alleles with C>N or both types of edits, while G>N editing efficiency was computed using alleles with G>N or both types of edits. Editing efficiencies inside and outside the −18 to −13 bp windows were calculated by only considering alleles that contained an edited base within −18 to −13 bp. Alleles containing no edits within the window were used to calculate editing efficiency outside the window.

### Analysis of editing window and base substitution spectrum or purity

The base substitution frequency table was used in this analysis. The reference base and position relative to the PAM were calculated for each base substitution. The position was calculated after reverse complementing target sites with sgRNAs targeting the bottom strand. Negative positions indicate the sgRNA is upstream of the PAM. The base substitution frequency at each base was summed for all substitutions at a frequency higher than 0.5%. This sum was divided by 100 to give the final aggregate editing value. To define whether a base position was edited, we calculated a cutoff (μ+2σ) based on the mean (μ) and standard deviation (σ) of aggregate editing at each position in the dCas9;MS2-AIDDead samples. The μ+2σ cutoffs were 0.189 for Transient and 0.211 for Integrated delivery systems. Therefore, 0.25 was used as a cutoff to call a base as edited for CRISPR-X (dCas9;MS2-AID*) samples. In the analysis of CRISPR-X, the target sites were separated based on whether the sgRNA targeted the top or bottom strand of the target site. We used a 0.5 cutoff to call a base edited for editors where the deaminase was directly fused to d/n/n’Cas9 because all target sites were combined for the analysis. The editing window was reported as the left- (least, including negative values) and right-most bases that exhibited editing above the cutoff. To calculate the editing spectrum of the base editors, we calculated the aggregate editing for all edits with the same reference base rather than an individual base position.

### dCas9 expression analysis using intracellular staining

Each condition was performed in three replicates with wild-type K562 cells. Lentiviral transduction and electroporation conditions were the same as the editing assay. For the electroporated samples, 3 days after electroporation, 4 × 10^6^ cells were collected from each replicate. For the transduced samples, 4 × 10^6^ cells were collected from each replicate 6 days after infection. Cells were fixed in Fix Buffer I (BD Biosciences 557870) at 37°C for 15 min and washed with PBS with 10% FBS. Cells were permeabilized with Perm Buffer III (BD Biosciences 558050) on ice for 30 min and washed again. They were incubated with primary anti-Cas9 (clone 7A9-3A3) antibody (ThermoFisher 61577, 1:1000 dilution) diluted in wash buffer at room temperature for 1 hr. They were then washed and incubated with anti-mouse secondary antibody conjugated to Alexa647 (ThermoFisher A-21235, 1:1000) diluted in wash buffer at 4°C for 1 hr in the dark. After incubation, cells were washed, resuspended in wash buffer, and measured via flow cytometry. For each sample, dCas9 positive and dCas9 negative cells were determined based on an uninfected/no electroporation control. The relative median dCas9 expression was calculated by normalizing the median dCas9 levels for dCas9 positive cells by the median dCas9 levels for the dCas9 negative cells.

### Measuring the distance between dCas9 termini and DNA bound by base editor

The 6vpc (19) structure was loaded into Pymol. To calculate the distance between dCas9 and the potential edited base, we selected the following atoms in the structure. For the N- and C-termini of dCas9, we selected the ɑ-carbon of the initial and final residue in the dCas9 structure, respectively. To represent the editing position, we selected the end of the nitrogenous base away from the phosphate backbone for the nucleotide at position -24 from the PAM on the targeted strand. The distance command in Pymol was used to calculate the distance between these positions.

### Motif enrichment score at edited cytidines or guanines

For each allele with a frequency above 0.5%, we extracted the reference sequence of the 7-mer centered on the edited base (−3 to 3 position). We produced the aggregate logo by summing each of these 7-mers at each relative position weighted by the frequency of the edited allele. The ES scores were calculated using a log-likelihood ratio based on the probability of the motif among the edited cytidines/guanines compared to a background distribution based on the 52% G/C content, which was used to generate the surrounding sequence. For example, in the WR<u>C</u> motif, the observed probability of the motif (P_observed_) = (frequency of A or T at position -2) × (frequency of G or A at position -1). The background probability (P_background_) is the same calculation based on the 52% GC content, which would be 0.48 × 0.5 = 0.24. The ES = Log_2_ (P_observed_ / P_background_). A positive ES score represents enrichment for the motif, while a negative value suggests the motif is depleted among the edited population.

### Quantifying base editor-induced diversification

The number of generated alleles was calculated considering alleles with a normalized frequency greater than 0.5%, excluding the unmodified reference allele. For each target site, the entropy was calculated as the sum of -*P* × ln (*P*), where *P* is the allele frequency for alleles with a frequency greater than 0.5%. This entropy calculation included the unmodified reference sequence as one of the alleles.

### Analysis of the number of edits per allele

We considered alleles with a normalized frequency above 0.5% and at least one base substitution in this calculation. The number of substitutions per allele was determined using the n_mutated field generated by CRISPResso2. The allele frequency for all such alleles was summed and grouped based on the number of mutations detected in the allele. This sum was normalized by the total editing frequency for each editor, and the proportion of the total edits for each editor was graphed.

### Quantification and Statistical Analysis

The graphs included in this study were generated using ggplot2 (20) or GraphPad PRISM. The logo graphs were plotted using ggseqlogo (21). Statistical analysis was performed using PRISM. Statistical analysis was two-sided (when applicable). Additional statistical test details are included in the figure legends.

## Results

### Design, construction, and analysis of paired sgRNA-target site library

To characterize the editing outcomes for genome editors for multiple target sites in parallel, we designed a lentiviral cassette that expressed a sgRNA next to a synthetic target site targeted by the sgRNA (**Figure 1A**). The sgRNA stem-loop contained two MS2 RNA hairpins (22) that can recruit cytidine deaminases fused to the MS2 coat protein (7). Oligonucleotides encoding sgRNA-target site pairs can be inserted into this cassette and delivered to mammalian cells. After adding a Cas9-based editor, the integrated cassette can be sequenced to determine the range of edits produced across all sgRNA-target site pairs for a population of cells.

**Figure 1.**
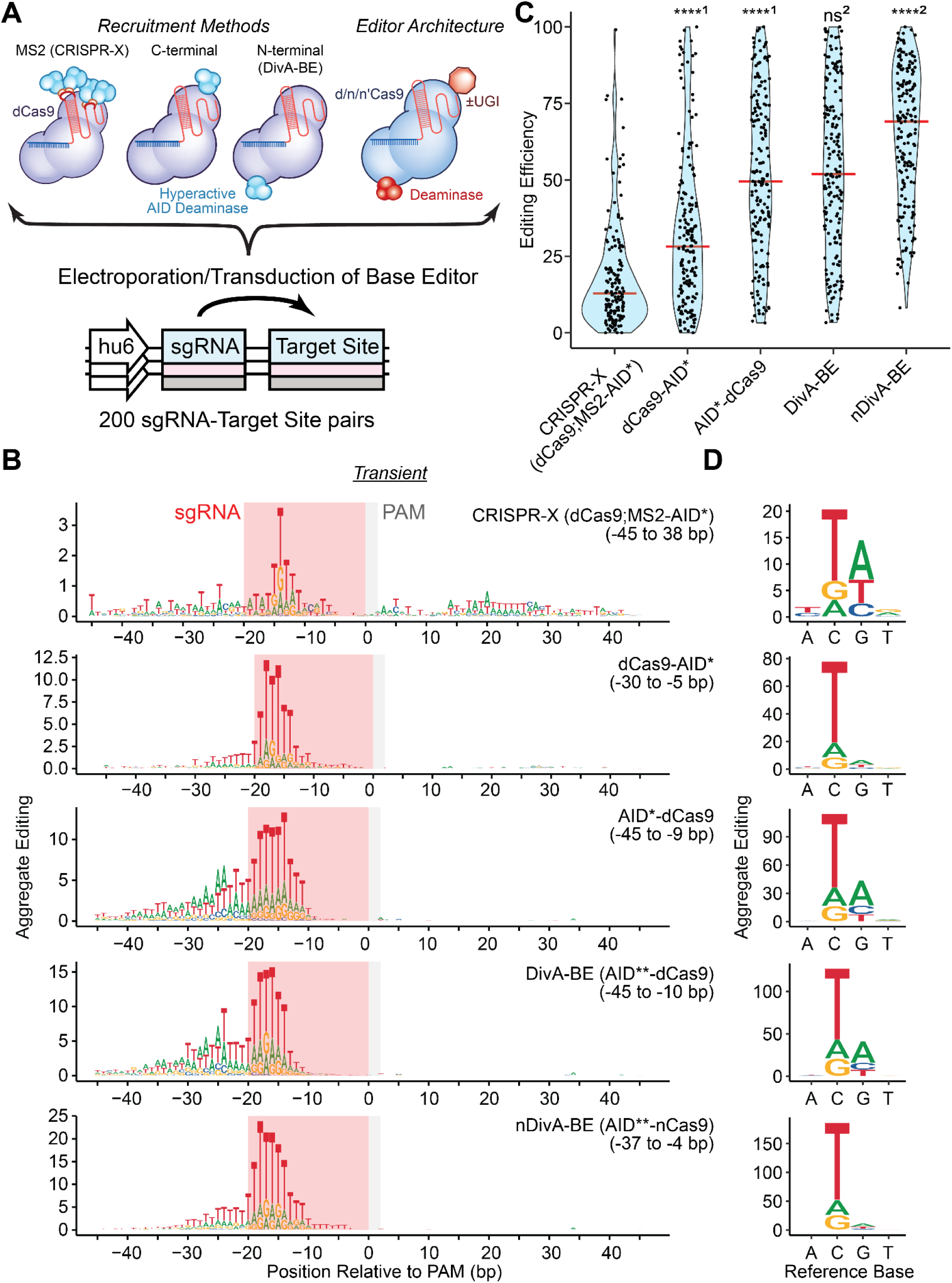
DivA-BE and AID variants fused to the N-terminus of dCas9 exhibited higher editing efficiency than C-terminal fusions or CRISPR-X. (A) Schematic for characterization of editing for 200 sgRNA-target site pairs. Base editors are introduced either via electroporation or lentiviral transduction to K562 cells containing a library of sgRNA-target site pairs. These editors contained variable recruitment methods for the deaminase and changes in editor architecture, including the version of Cas9, deaminase identity, and presence of a uracil glycosylase inhibitor (UGI). After editing, the sgRNA-target site amplicon is sequenced. (B) Aggregate editing logos of base substitutions produced by Transient CRISPR-X (dCas9;MS2-AID*), dCas9-AID*, AID*-dCas9, DivA-BE (AID**-dCas9), nDivA-BE (AID**-nCas9) base editors at each base position relative to the PAM. The bases shown indicate the resulting base substitution. The red-shaded boxes indicate the position of the protospacer sequence for sgRNAs. The gray-shaded boxes indicate the position of the NGG PAM. The left and right ends of editing windows relative to the PAM are indicated in parentheses. (C) Violin plots for editing efficiency of Transient CRISPR-X, dCas9-AID*, AID*-dCas9, DivA-BE, nDivA- BE base editors The red bar indicates the median editing efficiency. (Kruskal-Wallis test; ^1^ indicate comparisons to CRISPR-X and ^2^ indicate comparisons to AID*-dCas9; **** - p < 0.0001, ns - not significant) (D) Aggregate editing plot of editing spectrum of base substitutions generated by Transient CRISPR-X, dCas9-AID*, AID*-dCas9, DivA-BE, nDivA-BE base editors.

We designed 200 sgRNA-synthetic target site pairs to profile the editing activity of base editors in parallel (**Figure S1A**). sgRNAs without predicted targets in the human genome were selected to avoid any confounding growth effects from editing elsewhere in the genome. We reverse-complemented half of the resulting target site sequences to facilitate analysis of whether editing patterns were altered by which strand the sgRNA targeted. Ten negative control sgRNA-target site pairs were included, which should not be edited in the presence of a genome editor. During plasmid library construction, we inserted a 7-bp UMI to distinguish between oligonucleotide synthesis errors and base editor mutations in the downstream analysis to improve the detection of editing (23). PCR amplification of the resulting lentiviral cassette to attach sequencing adapters produces an amplicon compatible with paired-end high-throughput sequencing (**Figure S1B**).

The sgRNA-target site library was transduced into K562s, followed by selection with puromycin. To utilize the UMI to detect pre-existing mutations, we needed to determine the identity of all sgRNA-target site-UMI triplets in our cell population before editing was induced. The sequencing required to characterize all of the triplets for this population would not be feasible, so we bottlenecked the population to limit the diversity of sgRNA-target site-UMI triplets. The bottlenecked cells were expanded, and base editors were delivered transiently via electroporation (Transient) or integrated through lentiviral transduction (Integrated) (**Supplementary Table S3**). For Transient delivery, the expression vectors included a fluorescent marker (GFP or tagBFP), which was used to enrich transfected cells via FACS. For Integrated delivery, expression vectors contained antibiotic resistance markers (blasticidin or hygromycin) used to enrich transduced cells. After sorting or selection, genomic DNA was extracted from the uninfected parent cells and edited samples. We amplified and sequenced the sgRNA-target site cassette. The bottlenecking step, introduction of editors, and sequencing were performed in duplicate for all samples in the study.

We investigated multiple analysis pipelines to detect edits generated by base editors (**Figure S1C-D and Supplementary Text S2**) by analyzing the editing patterns of four samples from the Transient set: Parent (No electroporation), AID*-dCas9 (an N-terminal fusion of hyperactive AID (AID*) to dCas9), dCas9;MS2-AID* (CRISPR-X), and dCas9;MS2-AIDDead, a catalytically inactive version of AID. AID* was previously referred to as AID*Δ, which contains mutations (K10E, T82I, and E156G) that increase deaminase activity (24). The last three residues were removed to break the nuclear export signal (NES) and further improve activity. All AID variants in this study, including AIDDead, had no functional NES, so we dropped the Δ to simplify notation. The workflow of the final analysis pipeline is shown in **Figure S1E**.

### Transient delivery of editors produced higher editing efficiencies than Integrated delivery

We applied the pipeline to calculate editing efficiency for all sgRNA-target sites for all editors in the Transient and Integrated datasets (**Figure S2A**). This editing efficiency is the percentage of modified alleles produced by the base editor after normalization to the Parent sample that was not exposed to an editor. We observed low median editing efficiency for negative control sgRNA-target site pairs for the Transient (0.67%) and Integrated (1.37%) delivery. We calculated the correlation between allele frequencies for each target site and editors between the two replicates to determine whether our analysis was robust (**Figure S2B**). We observed a good correlation between the individual replicates for the Transient (Pearson=0.782) and Integrated (Pearson = 0.788) sets. These results demonstrate that our analysis method can detect editing robustly.

We observed a higher median editing efficiency for Transient On-Target sgRNA-target sites than Integrated ones (**Figure S2A**). However, the same editors were not tested in each dataset. Therefore, we calculated the median editing efficiency for the eight editors used with both delivery systems. We still observed an increase in median editing efficiency for Transient delivered editors (45.87%) compared to Integrated editors (30.71%). The normalized allele frequencies for each delivery system showed a modest correlation (Pearson = 0.518, **Figure S2C**), suggesting editing efficiencies across delivery systems were related. These results support that the Transient delivery of base editors generally produces higher editing efficiencies than Integrated delivery.

### Profiling by sgRNA-target site library captured the asymmetric editing pattern of CRISPR-X

To further validate our analysis pipeline, we tested whether our system could capture a known editing window of DCBEs. Previously, we defined the editing window of CRISPR-X (dCas9;MS2- AID*) (7) as -50 to 50 bp relative to the PAM. This window was dependent on the direction of transcription in the region. In our profiling assay, this dependency would result in different editing windows for sgRNAs that target the top strand compared to those that target the bottom strand. To determine the editing window for CRISPR-X, we calculated aggregate editing detected across all target sites for each base position in the CRISPR-X Transient and Integrated samples (**Figure S2D**). We divided the target sites based on whether the sgRNA targeted the top or bottom strand to test whether the editing windows differed between these two target site sets for CRISPR-X. We used a cutoff for calling a base as edited based on the dCas9;MS2-AIDDead sample and defined the editing window by the left- and right-most bases with aggregate editing above this cutoff. We observed an editing window of −45 to 38 bp for the Transient sample and −45 to 30 bp for the Integrated CRISPR-X sample. Both observed windows are narrower than the previously defined -50 to 50 bp window. This reduction could be due to a few reasons. First, we could not detect mutations in the −45 to -50 and 45 to 50 bp regions as these overlap primers used for sequencing preparation. Second, the editing window does not suggest that editing cannot occur outside the window, but the window indicates regions consistently edited across many sgRNA-target site pairs. Given the -50 to 50 bp window was derived from the editing pattern of two sgRNAs, our results may more accurately reflect areas of consistent editing for CRISPR-X.

We observed an asymmetric editing pattern of CRISPR-X for sgRNAs targeting the top or bottom strands (**Figure S2D**). For sgRNAs matching the bottom strand, we detected editing in the -27 to 14 bp window. However, for target sites where the sgRNA targeted the top strand, both delivery methods had an extended window from −45 to 30 bp. The observed strand-dependent editing windows are consistent with previous findings that the direction of transcription at the target site alters the editing window of CRISPR-X (7). These results demonstrate that our profiling assay can capture the editing patterns of Cas9-mediated base editors.

### AID* fused to the N-terminus of dCas9 had higher editing efficiency than C-terminal fusions and CRISPR-X

Multiple methods of recruiting the deaminase have been used in Cas9-mediated base editor systems (6, 25, 26). Yet, the effect of these recruitment methods on editing properties has not been studied over many target sequences. Thus, we analyzed the editing patterns for DCBEs that recruit AID* via an MS2 RNA aptamer system (CRISPR-X, dCas9;MS2-AID*), direct fusion to the C-terminus of dCas9 (dCas9-AID*), and a fusion to the N-terminus of dCas9 (AID*-dCas9) delivered transiently (**Figure 1B-C**). We observed a symmetric editing pattern for both direct fusions (**Figure 1B**), as all editing was detected upstream of the PAM. Given the symmetric editing for the fusions, we combined sgRNAs targeting the top and bottom strands in the calculation of aggregate editing. We found a reduced editing window for both fusions (−45 to -9 bp: AID*-dCas9, −30 to -5 bp: dCas9-AID*). Despite the reduced window, we saw an increase in median editing efficiency (**Figure 1C**) for AID*-dCas9 (3.88-fold) and dCas9-AID* (2.20-fold) compared to CRISPR-X.

For some assays studying the functional effects of SNVs, base editors are expressed constitutively after lentiviral transduction. Therefore, we investigated whether the method of AID* recruitment affected editing similarly for editors delivered after lentiviral integration (**Figure S3A- B**). Although we observed lower overall median editing efficiency for all Integrated editors (**Figure S2A**), we observed an increase in editing efficiency for CRISPR-X Integrated compared to Transient (18.14 vs 12.84). For the direct fusions, Integrated editors had a decrease in median editing efficiency compared to Transient editors. Surprisingly, the dCas9-AID* was the least efficient editor of the three Integrated editors, and we did not identify any positions with aggregate editing above the edited threshold. dCas9-AID* displayed a higher editing efficiency across all target sites when compared to the dCas9;MS2-AIDDead sample (p = 0.0216, Mann-Whitney test), suggesting dCas9-AID* was capable of editing alleles, albeit at low efficiency. AID*-dCas9 remained the most potent editor of the three recruitment methods tested with a similar editing window (−45 to -10 bp) in both delivery systems. These results show that the recruitment method of AID* influences the editing efficiency and target window in a delivery-dependent way.

The observed differences in editing efficiency between the recruitment and delivery methods could be due to dCas9 expression level variations. To determine if this was the case, we compared dCas9 levels using intracellular staining for dCas9, dCas9-AID*, and AID*-dCas9 expression vectors used in the Integrated and Transient datasets (**Figure S3C**). Each delivery method used different plasmid backbones to express dCas9. Transient delivery used a tagBFP vector, and the Integrated system used a vector containing a blasticidin resistance cassette. Both sets of plasmids were delivered by both delivery methods for this assay to determine whether the plasmids differentially expressed dCas9. We used dCas9 as a proxy for CRISPR-X, because dCas9 is required to target the editing site, and thus, its expression level is the limiting factor in editing. For both the Integrated and Transient delivery methods, dCas9 expression for dCas9- AID* and AID*-dCas9 were similar, yet AID*-dCas9 had higher editing efficiency (**Figure 1C and Figure S3B**). Regardless of the delivery method, dCas9 had higher expression levels than the fusion vectors, though this did not result in CRISPR-X being the most potent editor. Therefore, changes in editing efficiency between recruitment methods were not explained by dCas9 expression. We did not observe differences in dCas9 expression between plasmid backbones through either delivery method except for an increase in expression for the dCas9-tagBFP vector compared to its dCas9-blasticidin counterpart when delivered transiently. Surprisingly, we observed similar dCas9-AID* expression levels across delivery methods despite the differences in editing efficiency. Overall, dCas9 expression levels did not correlate with changes in editing efficiency across recruitment or delivery methods and do not explain the observed changes in editing efficiency.

An advantage of DCBEs is that they produce C>N mutations rather than only C>T, although the distribution of these edits has only been studied in low-throughput for a few target sites. We analyzed the editing spectrum of substitutions made by each base editor recruitment method **(Figure 1D and Figure S3D**). C>T was the most common substitution for all editors in this analysis. For Transient CRISPR-X, we observed a 67:16:18 distribution of C>T:A:G substitutions.

Both AID*-dCas9 (68:17:14) and dCas9-AID* (75:14:11) showed an increased preference for C>T mutations. For Integrated editors, we observed a similar preference for C>T edits, although C>G edits comprised a higher proportion (>30%) of the editing spectrum for Integrated CRISPR-X and dCas9-AID*. These results show that AID*-based DCBEs can effectively make C>N edits with a preference for C>T edits in the tested recruitment and delivery methods.

### DivA-BE and nDivA-BE, fusions of truncated hyperactive AID variant (AID**) to d/nCas9, increased editing efficiency

The AID*-dCas9 had the highest editing efficiency among the recruitment methods for either Transient or Integrated delivery. However, we sought to improve editing efficiency further. We tested two additional constructs using another hyperactive mutant AID**, which contained three additional mutations (S66T, L104I, and K160E) and truncated an additional 15 residues from the C-terminus of AID. AID** had previously been shown to be more mutagenic than AID* in an editing assay in bacteria (24). To determine if this deaminase could further improve editing efficiency, we fused AID** to dCas9, which we named the <u>Div</u>ersifying <u>A</u>ID fusion <u>B</u>ase <u>E</u>ditor (DivA-BE). Additionally, we fused AID** to nCas9 (nDivA-BE), which can increase base editing efficiency by biasing repair to use the deaminated non-target strand as the template in repair. For Transient delivery, we observed a small increase in median editing efficiency for DivA-BE compared to AID*- dCas9 (51.87 vs 49.88) (**Figure 1C**). However, nDivA-BE (69.06) exhibited an increase in editing efficiency over both dCas9 editors. In the Integrated delivery samples, we observed the same relative increase in editing efficiency (**Figure S3B**), although DivA-BE had a more substantial editing efficiency increase compared to AID*-dCas9 (27.01 vs 20.81).

The editing window for DivA-BE was similar to AID*-dCas9 (−45 to -10 bp: Transient, −39 to -11: Integrated) (**Figure 1B and Figure S2A**). Surprisingly, nDivA-BE had a shifted editing window towards the PAM for the Transient (−37 to −4 bp) and Integrated delivery (−36 to −3 bp). Despite the increased editing efficiency, AID**-nCas9 edited positions −45 to −38 less efficiently than the dCas9 fusions. Neither DivA-BE/nDivA-BE editors exhibited a change in C>T:A:G editing spectrum compared to AID*-dCas9, with 64-72% of cytidine edits resulting in C>T substitutions (**Figure 1D and Figure S3C**). These results show that DivA-BE and nDivA-BE have improved editing efficiency compared to AID*-dCas9.

### DivA-BE and AID*-dCas9 produced C>N and G>N mutations with shifted targeting windows

In addition to the expected C>N mutations, we observed the presence of G>N mutations for AID*- dCas9 and DivA-BE samples (**Figure 1D and Figure S3C**). We analyzed the editing efficiency of alleles that contain either C>N or G>N mutations (**Figure 2A and Figure S4A**) to determine whether the G>N mutations were present at a few target sites or were common across most editing sites. For both Transient editors and Integrated editors, we observed median editing efficiency greater than 10% for G>N alleles in the AID*-dCas9/DivA-BE samples, suggesting G>N editing occurred at most target sites tested in our assay. Furthermore, we noted that CRISPR-X, dCas9-AID*, and nDivA-BE samples had lower G>N editing efficiencies. This decrease in G>N editing could be due to reduced overall editing efficiency, so we computed the ratio of median G>N editing efficiency to median C>N editing efficiency. The relative G>N editing was lower in the dCas9-AID* (0.036) and nDivA-BE (0.065) samples, while CRISPR-X had higher relative G>N editing (0.605) compared to AID*-dCas9 (0.520), and DivA-BE (0.389) editors. We observed a similar relative depletion of G>N edits for Integrated nDivA-BE (0.040). The Integrated dCas9- AID* had a median G>N editing of 0, showing a G>N editing depletion. However, the median C>N editing was 0, so we could not compare the G>N:C>N ratio to other editors.

**Figure 2.**
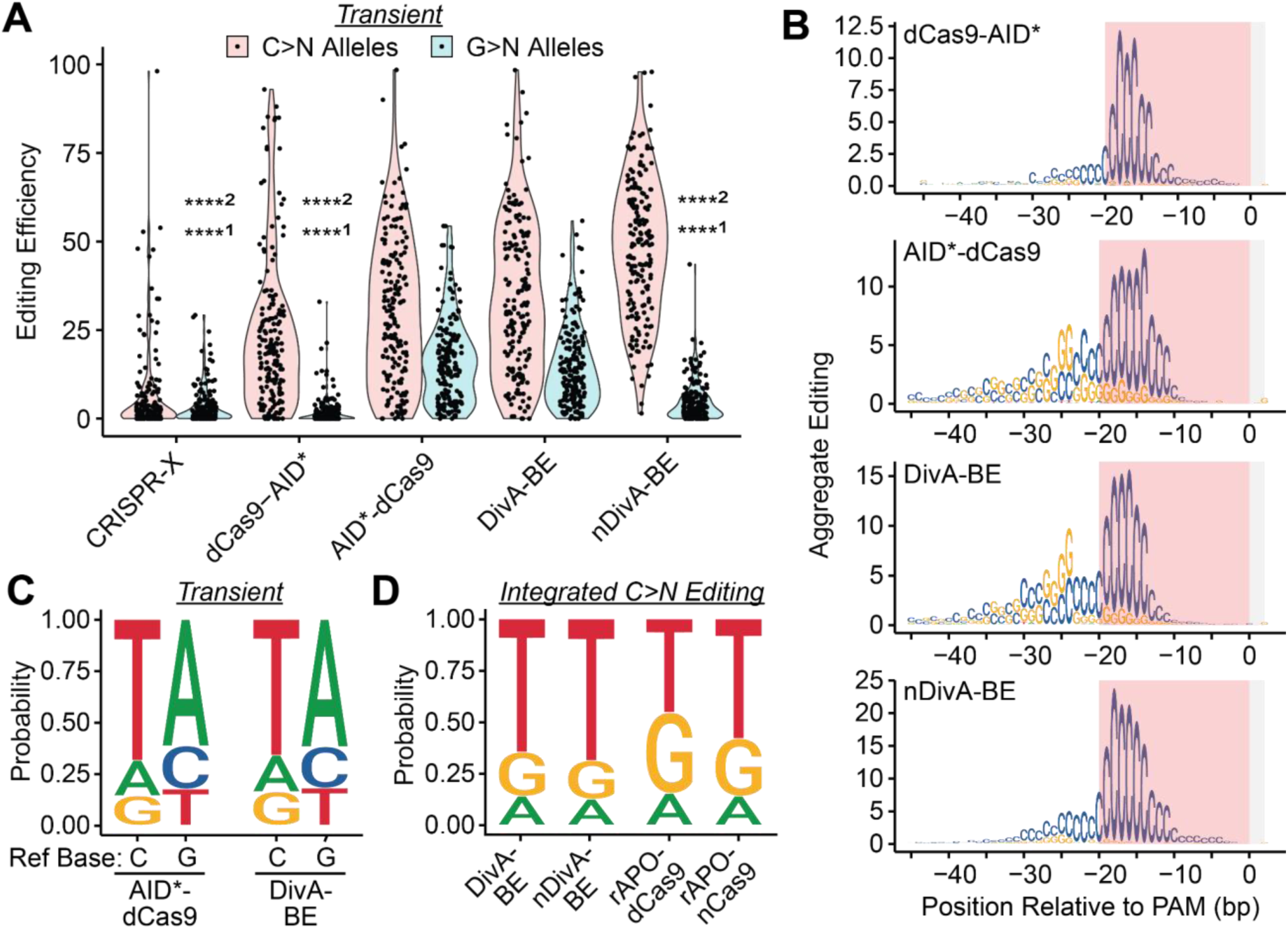
DivA-BE generated G>N mutations with a shifted window from C>N mutations. (A) Violin plots for editing efficiency of Transient CRISPR-X, dCas9-AID*, AID*-dCas9, DivA-BE, nDivA- BE base editors. Red violin plots represent the editing of alleles containing C>N mutations, and blue plots indicate efficiency for G>N mutations. (Kruskal-Wallis test; ^1^ indicate comparisons to G>N editing for AID*-dCas9 and ^2^ indicate comparisons to DivA-BE; **** - p < 0.0001) (B) Aggregate editing logos of reference base substitutions produced by Transient dCas9-AID*, AID*- dCas9, DivA-BE, and nDivA-BE base editors at each base position relative to the PAM. The bases shown indicate the reference base that was altered. The red-shaded boxes indicate the position of the protospacer sequence for sgRNAs. The gray-shaded boxes indicate the position of the NGG PAM. (C) Normalized editing spectra for C>N and G>N editing in AID*-dCas9 and DivA-BE samples for Transient delivery. (D) Normalized editing spectra for C>N editing in DivA-BE, nDivA-BE, rAPOBEC1-dCas9 (rAPO-dCas9), and rAPOBEC1-nCas9 (rAPO-nCas9) base editors for Integrated delivery.

nDivA-BE and dCas9-AID* exhibited lower levels of G>N editing and reduced editing efficiency outside the sgRNA target sequence (positions −45 to −20) compared to the AID*-dCas9/DivA-BE fusions (**Figure 1B and S3A**). We hypothesized that these two observations may be related. Therefore, we analyzed the reference bases being mutated and their positions for the Transient- delivered editors (**Figure 2B**). We observed a ∼20-25 bp G>N editing windows for the DivA-BE (- 35 to -14) and AID*-dCas9 (−37 to -11) and a narrower window for nDivA-BE (−19 to -27). Only base position -27 had G>N editing above our cutoff for Transient dCas9-AID*. We observed a similar trend for Integrated editors (**Figure S4B**) with G>N editing windows of -27 to -17 for AID*- dCas9 and −33 to −17 for DivA-BE. nDivA-BE and dCas9-AID* did not have any positions with aggregate G>N editing above our cutoff. For all of the editors where G>N editing was detected above our cutoff, the four positions with the highest G>N editing were in the -27 to -24 bp window, which is shifted from the −18 to −15 window for C>N editing. The shifted windows for G>N editing may explain the increased editing efficiency for AID*-dCas9/DivA-BE editors outside the sgRNA sequence window.

The most straightforward explanation for these G>N edits is from the deamination of the target strand. If these G>N edits were produced by AID-mediated deamination, we hypothesized that the distribution of edits (G>A:T:C) would be similar to those of C>T:A:G, since the same mechanism would repair them. We observed a similar distribution for G>A:T:C of 61:18:21 (AID*- dCas9) and 62:18:20 (DivA-BE), which aligns with the C>T:A:G distribution we observed for the same editors (**Figure 2C**). Similar concordance was present for the G>N and C>N editing spectrums for the Integrated samples (**Figure S4C**). Furthermore, deamination of the target strand is consistent with the depletion of G>N edits by fusing the nCas9, which nicks the target strand. This ssDNA break biases repair towards the non-target strand and disfavors repair of deaminations of the target strand. However, this model does not explain why dCas9-AID* has reduced G>N editing. We hypothesized that this may be due to the proximity of the dCas9’s termini to the G>N editing window. We analyzed the structure of an adenine base editor (19) to determine the distance of the N- and C-termini of dCas9 from the -24 bp nucleotide position, the closest positions with high levels of G>N editing for both AID*-dCas9 and DivA-BE. We analyzed this structure rather than cytidine base editor structures because it contained nucleotides that extended beyond the −20 bp window to where the maximal G>N edits occurred. The N-terminus was 4.63 nm away from the nucleotide, while the C-terminus of dCas9 was 8.31 nm from the -24 bp position. This increase in distance supports that the deaminase fused to the C-terminus may less efficiently interact with these positions of maximal G>N editing. Altogether, these results suggest that base editors with AID variants fused to the N-terminus of dCas9 produce G>N editing via deamination of the target strand.

### Nickase activity of Cas9 and deaminase identity altered relative G>N editing levels

Given the G>N edits were sensitive to nCas9, we hypothesized that introducing a Cas9 variant that nicks the non-target strand (n’Cas9) could increase the efficiency of G>N editing. Furthermore, we wanted to investigate whether base editors with other deaminases besides AID produced G>N edits. While the G>N editing window overlapped the editing window observed for other cytidine deaminases, AID-mediated base editors have extended editing windows compared to editors utilizing other deaminases (10), which may be necessary to facilitate G>N editing. To investigate these effects, we analyzed the editing patterns of DivA-BE, nDivA-BE, AID*-dCas9, and AID*-n’Cas9 delivered via Integration (**Figure S4A-B, and D**). We also analyzed the editing generated by rAPOBEC1 fused to dCas9, nCas9, and n’Cas9. We observed an increase in the ratio of G>N editing to C>N editing for AID*-n’Cas9 (0.821) compared to AID*-dCas9 (0.339) or DivA-BE (0.263). The G>N editing efficiency increase was slight (4.87 vs. 3.39) and not significant. This result suggests that the primary source of the relative G>N enrichment was from n’Cas9 reducing C>N editing (median: 5.93 vs. 13.68). For rAPOBEC1 samples, we observed low levels of G>N median editing (<2) for all fusions, similar to the levels observed for nDivA-BE (**Figure S4A**). When we analyzed G>N aggregate editing at each base position, we did not observe any positions above our cutoff in any of the rAPOBEC1 fusions. These results suggest that n’Cas9 can alter the relative G>N:C>N editing levels and that AID** makes G>N edits more efficiently than rAPOBEC1.

### Deaminase in the base editor altered the spectrum of base substitutions

For AID*/AID**, we found the editing spectrum to be ∼60:20:20 for C>T:A:G mutations. Previous studies have noted a difference in editing between deaminases (25, 27), but these experiments were performed in the background of a UGI, which biases the repair of the deaminated cytidines. To investigate whether deaminases affect the editing spectrum without the UGI, we compared the editing spectrum for cytidines edited by rAPOBEC1-d/nCas9 with DivA-BE/nDivA-BE samples (**Figure 2D**). Both deaminases preferentially introduced C>T edits. However, both rAPOBEC1 fusions showed an increased proportion of C>G edits (45:15:*<u>39</u>*–rAPOBEC1-dCas9 and 58:14:*<u>28</u>*–rAPOBEC1-nCas9). These results support the idea that deaminases can affect the editing spectrum of DCBEs.

### Editing window and purity of C>T programmable editors were affected by deaminase identity

Programmable C>T base editors have been used to study the effect of SNVs (28–30). In most of these studies, specific C>T mutations previously found in patient databases are installed, followed by phenotypic selection. The abundance of the sgRNA is the readout for enrichment/depletion, but this assumes that the mutation produces the expected mutation with high purity. Therefore, knowing the editing windows and spectrum for programmable editors is critical for their use in such applications. Thus, we compared the editing properties for three C>T editors (DivA-BE3, BE3 (rAPOBEC1), and PmCDA1-BE3 (cytidine deaminase from sea lamprey (4, 31)) in our profiling assay (**Figure 3A-B**). All these editors were in a BE3 configuration, consisting of a cytidine deaminase fused to nCas9 with a C-terminal fusion to a UGI. For Transient-delivered editors, DivA-BE3 was the most efficient editor (76.00%) and had the largest editing window (−3 to −45 bp). BE3 (−27 to −7) and PmCDA1-BE3 (−29 to −8) had similar editing windows, although BE3 had a higher median editing efficiency. We observed the same trends for the Integrated BE3 editors with BE3 (−27 to −7) and PmCDA1-BE3 (−25 to −10) exhibiting similar editing windows, while DivA-BE3 maintained the highest editing efficiency (**Figure S5A-B**). Despite having similar editing windows, we noted a rapid decrease in BE3 editing efficiency beyond the −18 to −13 bp window, suggesting a narrower editing window. Therefore, we quantified the editing for the Transient and Integrated BE3 editors inside and outside the −18 to −13 bp window (**Figure 3C and Figure S5C**). For BE3, the editor had ∼3-fold more editing in the narrow window (3.11-fold for Transient, 2.69-fold for Integrated), while AID** and PmCDA1 had 1.1-1.6-fold more editing in the narrow window, suggesting BE3 has a narrower editing window than the other two deaminases. These results demonstrate how deaminase identity alters the targeting window of C>T base editors.

**Figure 3.**
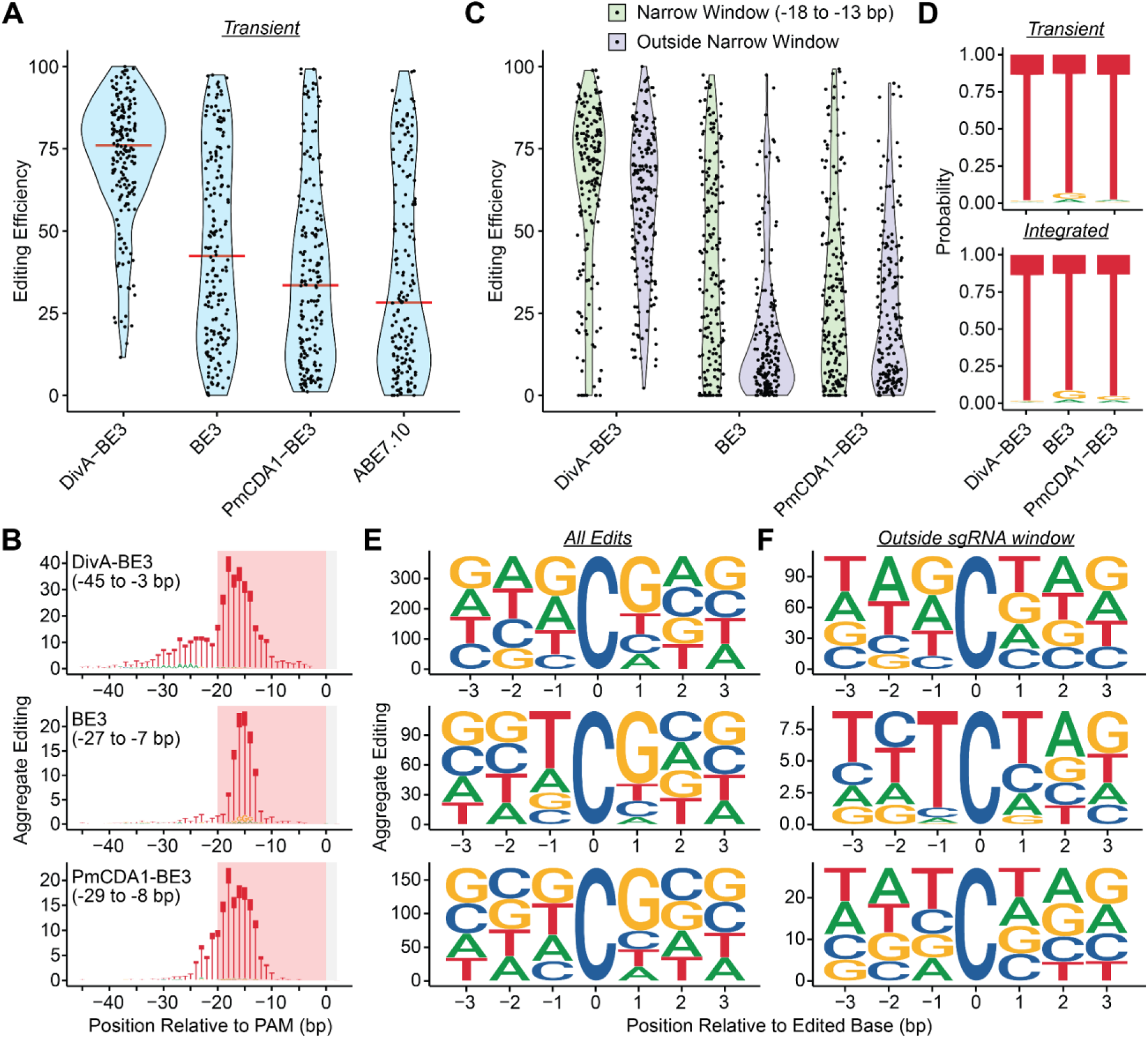
Editing window, spectra, and motif preferences for C>T editors were altered by deaminase identity. (A) Violin plots for editing efficiency of Transient DivA-BE3, BE3, PmCDA1-BE3, and ABE7.10 base editors. The red bar indicates the median editing efficiency. (B) Aggregate editing logos of base substitutions produced by Transient DivA-BE3, BE3, and PmCDA1- BE3 base editors at each base position relative to the PAM. The bases shown indicate the resulting base substitution. The red-shaded boxes indicate the position of the protospacer sequence for sgRNAs. The gray-shaded boxes indicate the position of the NGG PAM. (C) Violin plots for editing efficiency of Transient DivA-BE3, BE3, and PmCDA1-BE3 base editors. Green violin plots represent the efficiency of alleles containing edits in the narrow window (−18 to −13 bp), and the purple plots indicate the efficiency of alleles containing edits outside the narrow window. (D) Normalized editing spectra for C>N editing for Transient- and Integrated-delivered DivA-BE3, BE3, and PmCDA1-BE3 base editors. (E) Position-weight matrix for cytidines edited by Transient DivA-BE3, BE3, and PmCDA1-BE3. (F) Position-weight matrix for edited cytidines outside the sgRNA window (−20 to 0 bp) for Transient DivA- BE3, BE3, and PmCDA1-BE3.

Another critical property of C>T editors in assays assessing the function of variants is their ability to produce C>T editors and avoid other base substitutions. Adding a UGI impairs the activity of uracil-DNA glycosylases (32) needed to initiate base excision repair, which increases the proportion of C>T edits. Nonetheless, non-C>T mutations can occur, and these low levels of non-C>T edits can be enriched through phenotypic selection, leading to the improper annotation of the SNV causing the phenotype. Therefore, we analyzed the purity of the C>N edits produced by all three BE3 editors (**Figure 3D**). For the Transient delivered editors, we observed high C>T editing purity (>93%) for all editors. BE3 produced the lowest purity (93.23%) in comparison with DivA-BE3 (98.30%) and PmCDA1-BE3 (97.75%) editors. The Integrated editors maintained high C>T editing purity as well. BE3 (91.47%) had the lowest purity, while DivA-BE3 (98.29%) had the highest. For BE3, the impurities were most frequent within the narrow window (−18 to −13 bp), but the purity for cytidine edits at those bases was higher than those for the full editing window (93.77%:Transient and 92.23%:Integrated). This result suggests that the increased frequency of non-C>T edits was due to higher deamination activity at these positions rather than any differences in the repair of those base positions.

Adenine base editors (ABEs) have also been used to assess the function of SNVs (33). Therefore, we profiled the ABE editor ABE7.10, which was delivered transiently. This editor consists of an engineered dimer of TadA, a bacterial adenosine deaminase, fused to nCas9 (34). The deaminase converts an adenosine to an inosine, which leads to an A>G conversion during DNA replication. The median editing efficiency for the ABE7.10 (28.26) was lower than that of the programmable CBEs (**Figure 3A**), but it had the narrowest editing window (−18 to −10 bp) (**Figure S3D**). The editing purity for ABE7.10 was similar to BE3 editors at 95.67% A>G purity. Although TadA’s primary function is deaminating adenines, we found a low frequency of cytidines being mutated at -15 to -13 bp, the three most efficiently edited positions for the ABE (**Figure S3D**). These findings are consistent with other reports that the engineered TadA enzyme can deaminate cytidines, albeit less efficiently than adenines (35). These findings support that ABE7.10 is an efficient A>G editor, but cytidines in that window may be subject to low-frequency mutagenesis.

### Cytidine deaminase motif preferences were enhanced outside the sgRNA-targeting window

Cytidine deaminases edit cytidines in specific motifs more efficiently, and the motif preference differs between deaminases. In their native function, the deaminases may interact with other proteins to target a cytidine for deamination, but the targeting preference could be altered when the deaminase is recruited by d/nCas9. Therefore, we analyzed the motifs of the cytidines edited by the three BE3 editors (**Figure 3E and Figure S5E**). Previous studies have found that AID preferentially edits a WR<u>C</u> editing motif (36), while the rAPOBEC1 preferentially edits a T<u>C</u> motif (8). To test the preference for these motifs, we calculated a motif enrichment score (ES) for each motif based on the observed editing. For this score, a positive value represents an enrichment for that motif, while a negative score represents depletion. For DivA-BE3, we observed a positive enrichment for the ES_WR<u>C</u>_ of 0.603 for the Transient editor and 0.558 for the Integrated editor. Other reports have found a WR<u>C</u>Y motif for AID (37, 38), but we found that including this extra base in the motif lowered the ES_WR<u>C</u>Y_ score (0.116:Transient and 0.063:Integrated). For BE3, we observed enrichment of the T<u>C</u> motif in both Transient (ES_TC_ = 1.078) and Integrated (ES_TC_ = 1.109) samples.

For all three BE3 base editors, we observed maximal editing within the −20 and 0 bp positions. These positions undergo more frequent deamination due to the proximity of the deaminase to this region due to its fusion to nCas9, independent of the deaminase’s motif preference. Consequently, edits outside this window may exhibit an enhanced motif preference, as these edits are less favored by the fusion of the deaminase to nCas9. To test this possibility, we performed the motif enrichment analysis of edited cytidines but only included edits outside the −20 to 0 bp region (**Figure 3F and Figure S5F**). We observed an increase in ESs for the known editing motifs. For DivA-BE3, we found ES_WR<u>C</u>_ to be 0.943 and 0.958 for Transient and Integrated editors, respectively. Enrichment for T<u>C</u> was increased for BE3 with ES_T<u>C</u>_ = 1.827 (Transient) and 1.902 (Integrated).

For PmCDA1, a strong preference for AT<u>C</u> was identified when the enzyme was overexpressed in yeast (39). We calculated an enrichment score and observed a weak enrichment for the AT<u>C</u> motif (ES_AT<u>C</u>_ = 0.073:Transient and 0.236:Integrated). When this analysis was applied to cytidine edits outside the −20 to 0 bp window, we observed an increase in ES_AT<u>C</u>_ scores (1.013 and 1.427). These results show that the motif preference for cytidine base editors is enhanced outside the sgRNA targeting window.

### G>N editing has similar motif preferences as C>N editing for AID*/**-dCas9 fusions

Given the enhanced enrichment of ES scores for cytidines edited outside the −20 to 0 bp window, we examined whether G>N sites exhibited a similar enhancement of motif preference. We calculated ES scores for AID*-dCas9/DivA-BE samples for the WR<u>C</u> and WR<u>C</u>Y motifs for cytidines and <u>G</u>YS and R<u>G</u>YS motifs for guanines (**Figure S5G**). We performed this analysis for edits across the full window and those outside the sgRNA window. We observed enrichment for the WR<u>C</u> motif and enhanced enrichment for edits outside the sgRNA window, similar to what was observed for the BE3 editors. Interestingly, the WR<u>C</u>Y motif was less enriched than the WR<u>C</u> motif for C>N mutations across the full window, but ES_WR<u>C</u>Y_ was greater than ES_WR<u>C</u>_ for C>N mutations outside the sgRNA window, exhibiting increased enrichment of the extended motif. For G>N mutations, the <u>G</u>YS motif showed higher enrichment than WR<u>C</u> across the full editing window. However, restricting the analysis to mutations outside the sgRNA window did not substantially enhance the motif preference. The lack of enhanced motif preferences was unsurprising since most G>N mutations occur outside the sgRNA window. The extended R<u>G</u>YS motif increased the ES for edited guanines outside the sgRNA window, although the magnitude of this enhancement was less than the WR<u>C</u>Y enrichment over the same window. These results suggest that G>N editing has similar motif preferences to C>N editing for AID*-dCas9/DivA-BE fusions.

### The introduction of a uracil glycosylase inhibitor reduced the indel efficiency of cytidine base editors

An advantage of base editors over other genome editing strategies for investigating the function of protein-coding variants is avoiding double-strand break intermediates. Bypassing this intermediate reduces the toxicity of the genome editor and avoids the production of insertions/deletions (indels), which can knock out the protein function via a frameshift rather than install the variant of interest. Therefore, we compared the frequency of alleles containing indels for the editors in these studies (**Figure S6A-B**). We observed low median indel editing efficiency (<10%) for all editors except for nDivA-BE. The proposed mechanism for cytidine base editors producing indels is the excision of the deaminated cytidine to form an abasic site that an endonuclease can process to create a ssDNA break. The ssDNA break combined with other spontaneous nicks/ssDNA breaks or those generated by additional deamination can form double-strand breaks, which result in indels. Therefore, we hypothesized that changes in indel efficiency could be due to changes in deamination efficiency, which would result in increased editing efficiency. We calculated the correlation between the median indel editing efficiency and overall editing efficiency for all cytidine base editors tested in our assay (**Figure S6C**). We observed a low to moderate correlation (Pearson_All_ = 0.509-Transient and 0.381-Integrated). However, we noted that the three BE3 editors, which contain a UGI that inhibits the excision of the deaminated cytidine, exhibited low indel efficiency relative to their efficiency. Therefore, we calculated the Pearson correlation excluding the DivA-BE3, BE3, and PmCDA1-BE3 samples and observed a high correlation between indel and editing efficiency among the other editors (Pearson_NoUGI_ = 0.995-Transient and 0.830-Integrated). These data support the model that the base excision step is critical for producing indels induced by cytidine base editors.

### Efficiency and window size affected the diversity and number of edits produced by DCBEs

The ability of DCBEs to generate alleles that can be interrogated for phenotypic effects is critical for their usage in assays studying SNVs. We assessed the ability of the editors in the study to diversify their target sites using two metrics. The first was the number of alleles generated by a single sgRNA (**Figure 4A and Figure S7A**). The second was the entropy generated by each sgRNA, as this measurement would consider the frequency of the generated alleles (**Figure 4B and Figure S7B**). For both Transient and Integrated editors, nDivA-BE produced the highest median number of alleles (30–Transient and 35–Integrated) and the highest median entropy (3.224–Transient and 2.562–Integrated). These differences in entropy and number of alleles could be due to changes in editing frequency. To determine the contribution of editing efficiency to these properties, we calculated the correlation between median editing efficiency and median entropy and number of alleles (**Figure S7C**). We observed a strong correlation between median editing efficiency and the number of alleles and entropy for each editor (Pearson_NumAlleles_ = 0.764 and Pearson_Entropy_ = 0.942). However, efficiency is not the only determinant of diversifying ability, as DivA-BE3, our most efficient editor, was not the top diversifier by either metric.

**Figure 4.**
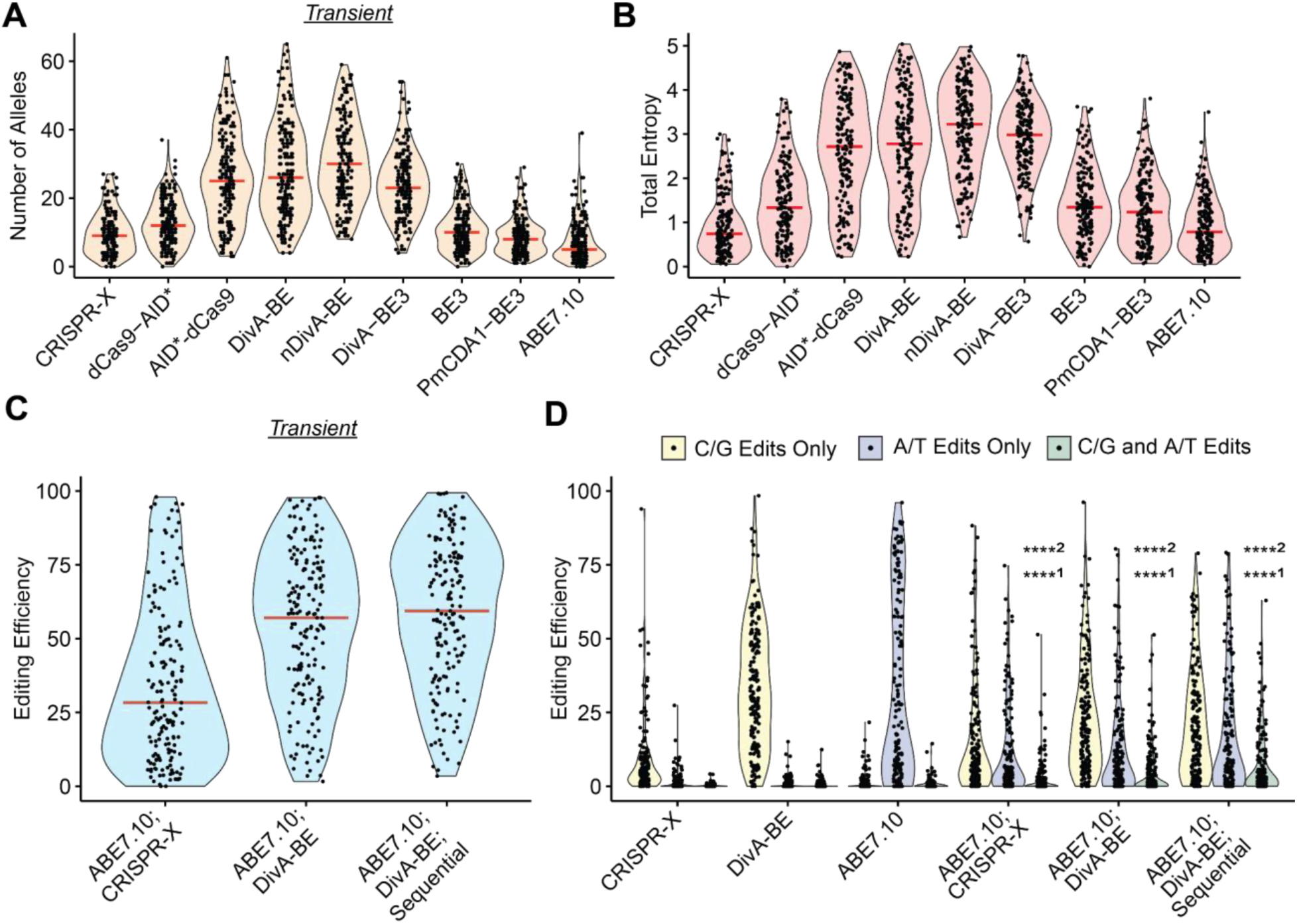
Cytidine base editors diversified target sites alone or in combination with adenine base editors. (A) Violin plots for the number of alleles generated by Transient base editors. Red bars indicate the median number of alleles generated. (B) Violin plots for the total entropy generated by Transient base editors. Red bars indicate the median entropy generated. (C) Violin plots for editing efficiency of combined of cytidine and adenine base editors: ABE7.10;CRISPR-X, ABE7.10;DivA-BE, and ABE7.10;DivA-BE;Sequential. Red bars indicate the median editing efficiency. (D) Violin plots for editing efficiency alleles containing C/G edits only, A/T edits only, or both types of edits for combinations of cytidine and adenine base editors: ABE7.10;CRISPR-X, ABE7.10;DivA-BE, and ABE7.10;DivA-BE;Sequential. (Kruskal-Wallis test; ^1^ indicate comparisons to editing efficiency of alleles containing both edits for the corresponding cytidine editor, and ^2^ indicate comparisons to ABE7.10; **** - p < 0.0001)

Bystander mutations where other bases are mutated besides the desired mutation can confound the interpretation of which SNV is causal in high-throughput base editor variant screens. Therefore, we analyzed the alleles generated by the editors to determine the number of mutations on each allele (**Figure S7D-E**). For the Transient delivered editors, alleles with a single mutation had a plurality in all conditions except for nDivA-BE, DivA-BE3, and PmCDA1-BE3. For nDivA- BE and PmCDA1-BE3, alleles containing two mutations had plurality (29.2% and 21.9%, respectively). For DivA-BE3, alleles with three mutations had plurality (22.5%), although this was nearly equivalent to the proportion of alleles with two mutations (22.0%).

These alleles with multiple edits could be due to repeated deamination events or the processivity of the deaminase. Only CRISPR-X, BE3, and ABE7.10 had > 50% of edited alleles containing a single mutation. BE3 and ABE7.10 had the narrowest editing windows of all the editors analyzed, suggesting that the processivity of the enzymes may play a critical role in controlling the number of edits on an allele. However, this is unlikely to be the sole determinant as CRISPR-X had the largest editing window of any of the Transient delivered editors investigated. To further distinguish the effect of multiple deamination events, we performed the same analysis on the number of mutations per allele for the Integrated editors (**Figure S7E**). Given these editors were constitutively expressed, they have a greater chance of deaminating a site repeatedly since a single mutation may not disrupt Cas9 re-targeting the target site, especially if the mutation is outside the sgRNA binding site. Other than dCas9-AID*, we did not observe changes in the distribution of the alleles based on the number of mutations compared to the results for the Transient editors. The change for dCas9-AID* was not surprising since the Integrated sample exhibited a reduction in editing efficiency (**Figure 1C and Figure S3B**). Similar to the Transient samples, DivA-BE3, nDivA-BE, and PmCDA1-BE3 were the only Integrated editors where alleles containing more than one mutation had a plurality. These results suggest that the editing window is one critical factor in determining the number of mutations per allele.

### Combinations of ABE and AID-based editors edited alleles in parallel and sequentially

Our previous analysis investigated the ability of either a CBE or ABE to work individually. Combining these two base editors in a single assay could generate additional diversity that can be interrogated for phenotypic effect. Thus, we transiently delivered ABE7.10 combined with either MS2-AID* (ABE7.10;CRISPR-X) or DivA-BE (ABE7.10;DivA-BE). Additionally, we performed sequential editing by first delivering ABE7.10 and a subsequent round of electroporation to deliver DivA-BE (ABE7.10;DivA-BE;Sequential). Both samples with DivA-BE had a higher median editing efficiency (**Figure 4C**) than the ABE7.10;CRISPR-X sample. Sequential delivery of ABE7.10 and DivA-BE had the highest editing efficiency (59.98). We assessed the ability of these combined editors to diversify their target sites by calculating the number of alleles (**Figure S7F**) and entropy (**Figure S7G**) generated by each combination. We observed the highest median number of alleles (31) and entropy (2.90) for the sequential editor combination. These values were similar to those of nDivA-BE, but these were increases over DivA-BE or ABE7.10. For each editor combination, the ABE and CBE could be editing alleles independently, such that no allele would contain both a mutated cytidine and adenine. To determine whether co-editing occurred, we evaluated the frequency of alleles that contained only C/G edits, A/T edits, or both (**Figure 4D**). For alleles containing both types of edits, we observed an increase in mean editing efficiency for all three combination editors, although only the combinations with DivA-BE increased the median. ABE7.10;DivA-BE;Sequential maintained the highest mean (7.55) and median (2.61) among the combinations tested. These results show that the combination of these editors can make multiple edits to a single allele and that sequential delivery of these editors leads to a modest increase in editing efficiency and diversification.

## Discussion

High-throughput assays using programmable and diversifying base editors have great potential to identify SNVs that profoundly impact cellular/gene function or disease progression. Additionally, these tools can identify loss- and gain-of-function mutants that can uncover the molecular mechanisms for a gene of interest. Characterizing the editing properties of genome editors is critical for assessing their suitability for these assays. Our results show the value of performing these characterizations across many targets as the context (e.g., surrounding sequence) can alter the editing outcomes. For example, our results defined a narrower editing window for the CRISPR-X editor, which was originally defined from only two sgRNAs. While this study marks the most comprehensive characterization of diversifying base editors to date, programamble base editors have been profiled for thousands of sgRNA-target site pairs (10, 11), enabling the application of machine learning algorithms to predict editing outcomes. Studies at this scale would be valuable for diversifying base editors, although the smaller size library used in this screen may be more feasible when studying many base editor configurations to prioritize editors for more intensive characterization.

Our studies established the DivA-BE editor as a more potent diversifying editor compared to CRISPR-X. This new editor had ∼4-fold increase in editing efficiency. Fusing the deaminase to the N-terminus of dCas9 drastically increased editing efficiency for either Transient or Integrated delivery methods. Fusing AID* to the C-terminus of dCas9 (dCas9-AID*) did increase editing efficiency when delivered transiently, although not to the level of the N-terminal fusions. However, surprisingly, this increase in editing efficiency for the C-terminal fusion was absent for Integrated delivery. The relative expression of the editors did not explain differences in editing efficiency between the delivery and recruitment methods. For example, dCas9-AID* and AID*-dCas9 had similar expression levels for both integrated and Transient delivery, but AID*-dCas9 had higher editing efficiency. Transient delivery is associated with higher expression levels than lentiviral constructs, which may explain the decrease in editing efficiency for dCas9-AID* with Integrated delivery. However, we found similar expression levels for dCas9-AID* in both delivery methods, although this does not rule out that higher expression levels were present at an earlier time point before dCas9 expression was assayed. While our results suggest that differences in editing efficiency between recruitment methods are not due to expression levels alone, delivering the editors as mRNA or ribonucleoproteins may be necessary to control dosage and definitively remove the effects of expression.

Despite the increase in editing efficiency, the direct fusions of AID variants (AID*/AID**) to dCas9 had reduced editing windows compared to CRISPR-X. The difference in editing windows between fusions and MS2 recruitment may not extend beyond AID variants, as recent work showed that recruitment of rAPOBEC1 variants through an RNA aptamer system did not have an extended editing window compared to the direct fusions of rAPOBEC1 (40). Both AID*-dCas9 and DivA-BE exhibited targeting windows of > 20 bp, wider than those generated by rAPOBEC1 and PmCDA1 that were tested in this study. The expanded editing window for base editors employing AID can be advantageous for targeting PAM deserts in studies of SNVs that affect gene function. This limitation may be mitigated by fusing a deaminase to CRISPR/Cas proteins with altered PAM preferences (41). Still, the increased editing window will reduce the number of sgRNAs required to target large genomic regions, making screens targeting these areas more tractable.

The method of fusing AID*/** to dCas9 affected the base substitution spectrum of diversifying base editors. The N-terminal fusions efficiently installed G>N edits and canonical C>N edits. These edits are initiated by deamination of the target (G>N) and non-target strand (C>N), as the G>N edits were depleted when the deaminase was fused to nCas9, which biases repair to the non-target strand. However, surprisingly, using the n’Cas9, which biases repair to use the target strand, produced a slight, non-significant increase in G>N editing. Recent reports have found that n’Cas9 with engineered AID variants increased G>N editing (12), although these editors contained a UGI, unlike the editors tested in our study. Notably, these G>N edits were depleted for dCas9-AID*. We hypothesized that this may be due to the inability of the deaminase to interact with the region of G>N editing when fused to the C-terminus of dCas9. Structural studies of base editors complexed with their DNA target show that the C-terminus of dCas9 is further from the nucleotide positions of maximum G>N editing than the N-terminus of dCas9. Furthermore, C>N editing at those positions is reduced for dCas9-AID* compared to AID*-dCas9, further supporting our hypothesis that the deaminase’s interactions with these positions are restricted. Investigating the effect of extending the linker length between the C-terminus of dCas9 and AID* on G>N editing levels may uncover strategies to modulate the levels of target strand deamination.

These additional G>N edits have implications for the applications of base editors in therapeutic and research applications. For DCBEs, G>N editing may allow the installation of additional alleles that can be interrogated for phenotypic effect. Furthermore, the possibility of G>N editing is a consideration for unwanted bystander mutations in therapeutic applications. Although we did not detect significant G>N editing for rAPOBEC1 base editors, the existence of G>N editing for AID*- dCas9 and DivA-BE suggests a ssDNA substrate must be present for AID to deaminate the target strand. The ssDNA substrate may be targetable by other base editors or endogenous deaminases. While our data finds a low frequency of these edits when used with nCas9, they should be carefully considered as a source of bystander edits in base editor applications.

Our study revealed how deaminases alter the base editing properties for programmable C>T editors. DivA-BE3 had an extended editing window of over 40 bp, while rAPOBEC1-BE3 and PmCDA1-BE3 had windows of ∼20 bp. This extended window aligns with studies showing that AID is a more processive deaminase (42, 43), consistent with its function in somatic hypermutation, where it deaminates the 1-2 kb Ig region. The distribution of the editing within the window also differs between deaminases. Although rAPOBEC1- and PmCDA1-BE3 have similar editing windows, rAPOBEC1-BE3 more efficiently edited the −18 to −13 bp, representing a more narrow editing window for that deaminase. Our study reported extended editing windows compared to those previously documented for rAPOBEC1 and PmCDA1. This difference may be due to having a lower threshold for calling a base edited. However, these extended editing windows are more applicable to high-throughput assays as they more accurately represent sources of low-frequency mutations that can be enriched by phenotypic selection, which may confound the annotation of the causal mutation.

The deaminase utilized in a base editor can alter the editing spectrum for diversifying and programmable base editors. For the BE3 editors, DivA-BE3 was the most pure editor for producing C>T edits (>98%), while rAPOBEC1 was consistently the lowest (91-93%). These are consistent with previous findings, and newer versions of rAPOBEC1-based editors like BE4 include two UGIs to improve C>T purity (27). Without a UGI, there were notable differences in editing spectrums as rAPOBEC1-d/nCas9 fusions had a higher frequency of C>G mutations than DivA-BE/nDivA-BE editors. This finding may explain an observed increase of C>T and C>G mutations throughout the genome in iPSCs expressing rAPOBEC1 base editors (44). Furthermore, variants of rAPOBEC1 have been developed with enhanced C>G editing (45, 46). These results suggest that rAPOBEC1 may engage factors that alter the repair of the deaminated cytidines. One potential factor involved in the repair of rAPOBEC1 base editing could be the REV1 translesion polymerase. *REV1* is responsible for inserting a guanine opposite of the cytidine (47), and its knockdown was found to alter C>G editing in a CRISPRi screen (46).

Base editors have been used extensively to annotate the phenotypic effects of SNVs. These assays are performed in one of two ways: installing known variants using programmable editors or tiling regions to discover mutations. For installing specific C>T variants, we suggest that base editors utilizing rAPOBEC1, or other editors with narrow editing windows, are preferable to AID** and PmCDA1 editors despite DivA-BE3 being more efficient. DivA-BE3 produced a higher frequency of alleles containing more than one mutation, which makes interpreting the causal mutation difficult in these experiments. We recommend DivA-BEs for variant discovery due to the increased window size and diversification potential. While nDivA-BE creates the most diversity (as measured by the number of alleles or entropy), it comes with tradeoffs. Similar to DivA-BE3, nDivA-BE most commonly produces alleles with multiple mutations. Additionally, nDivA-BE produced the highest frequency of indels, which may produce unwanted knockouts when editing protein-coding regions. Therefore, we recommend the DivA-BE as a versatile editor for these discovery assays in large genomic regions.

In summary, we developed an assay to profile the activity of diversifying base editors and define their editing patterns for ∼200 target sites. These studies reveal several properties that regulate base editing, including mutations induced by the deamination of the target strand. These findings reveal strategies to alter base editing properties by deaminase recruitment and potential sources of unwanted bystander mutations. The results of these studies will assist in selecting, applying, and interpreting the outcomes of base editor experiments assessing the function of point mutations and clinically identified SNVs.

## Data Availability

The sequencing datasets generated are deposited at the Sequencing Read Archive. Plasmids under Bioproject PRJNA1180481 (https://www.ncbi.nlm.nih.gov/bioproject/PRJNA1180481). Plasmids used in this study will be deposited at Addgene and are available upon request. The scripts used in this analysis are available on GitHub (https://github.com/HessLabUW/BECharacterization).

## Supplementary Data Statement

Supplementary Data is available on *bioRxiv*.

## Acknowledgments

We thank Michael Bassik for providing reagents to construct the plasmid library. We thank members of the Hess lab for helpful discussions. We thank Catherine Fox for her valuable feedback on the manuscript.

## Author Contributions

The research was conceived by G.T.H. C.I.S., A.L., A., A.T., and G.T.H. performed experiments. N.S.A. and G.T.H. performed data analyses with critical input from J.T. and S.B.M. G.T.H. and C.I.S. wrote the paper with help from all the authors.

## Supplementary Data

**Supplementary Figures S1-S7 Supplementary Text S1-S2**

**Supplementary Table 1: List of synthesized oligonucleotides**

**Supplementary Table 2: List of oligonucleotides and plasmids used in the study Supplementary Table 3: List of editing combinations used in the study**

