## Supplementary Information for "Towards optimizing diversifying base editors for high-throughput studies of single- nucleotide variants"

**Supplementary Data:**

**Supplementary Figures S1-S7**

**Supplementary Text S1-S2**

**Supplementary Table 1: List of synthesized oligonucleotides**

**Supplementary Table 2: List of oligonucleotides and plasmids used in the study**

**Supplementary Table 3: List of editing combinations used in the study**

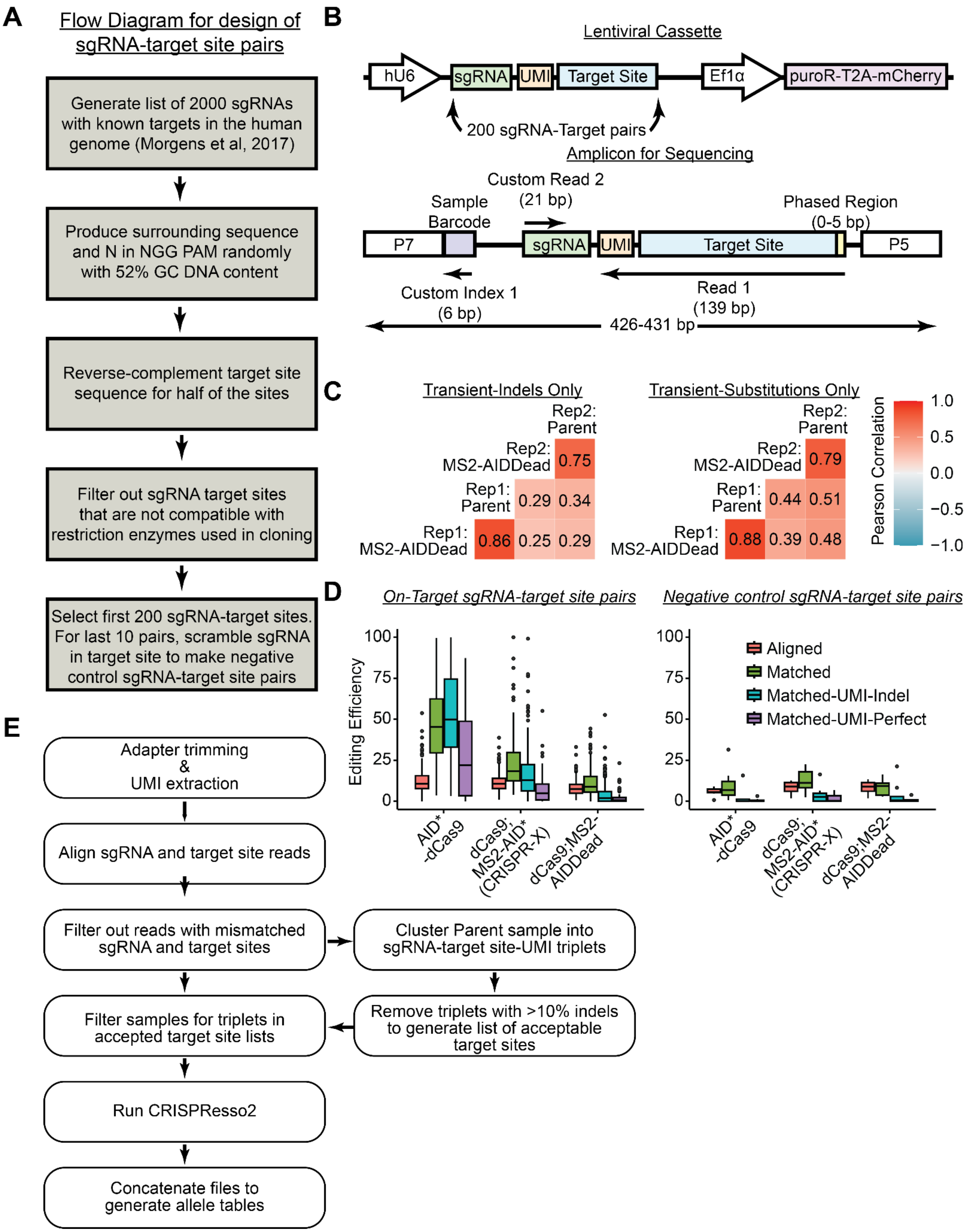


**Supplementary Figure S1. Using sgRNA and UMI in analysis improved the detection of base editing efficiency in sgRNA-target site profiling assay**

(A) Workflow diagram for designing 200 sgRNA-target site pairs

(B) Schematic for the design of lentiviral cassette and the amplicon that is compatible with paired-end sequencing
(C) Pearson correlation between allele frequencies for alleles that contain indels only or substitutions only between Parent and MS2-AIDDead samples for both replicates
(D) Boxplots of editing efficiency for AID*-dCas9, dCas9;MS2-AID* (CRISPR-X), dCas9;MS2-AIDDead samples for Aligned, Matched, Matched-UMI-Indel, and Matched-UMI-Perfect analysis methods. Boxplots are shown for editing efficiencies for On-Target sgRNA-target site pairs and negative control sgRNA-target site pairs. The whiskers represent the 1.5x of the interquartile range. The circles indicate outliers.
(E) Workflow of filtering and analysis pipeline to generate allele tables for all editing samples

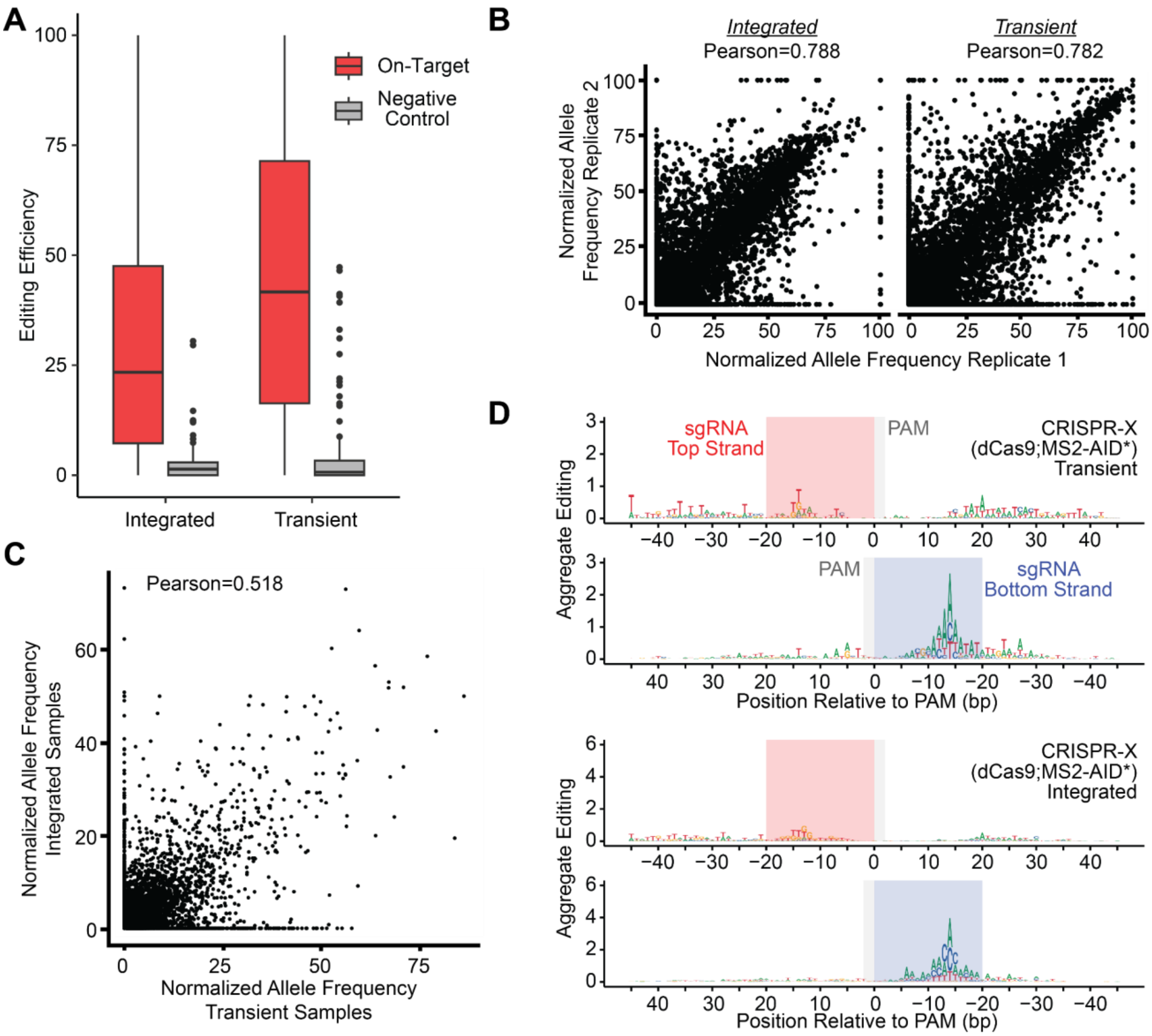


**Supplementary Figure S2. Paired sgRNA-target site library robustly identified base editing patterns**

(A) Boxplots of editing efficiency for all sgRNA-target site pairs among all editors. The whiskers represent the 1.5x of the interquartile range. The circles indicate outliers.

(B) Scatterplots of normalized allele frequencies between Replicate 1 and Replicate 2 for Integrated and Transient delivered editors. Pearson correlation between replicates is indicated above the scatterplots.

(C) Scatterplot of normalized allele frequencies for editors common to both Integrated and Transient datasets. Pearson correlation between editing produced by each delivery method is indicated on the graph.

(D) Aggregate editing logos of base substitutions produced by CRISPR-X editor with Transient or Integrated delivery. The bases shown indicate the resulting base substitution. The red- and blue-shaded boxes indicate the position of the protospacer sequence for sgRNAs targeting the top and bottom strands, respectively. The gray-shaded boxes indicate the position of the NGG PAM.

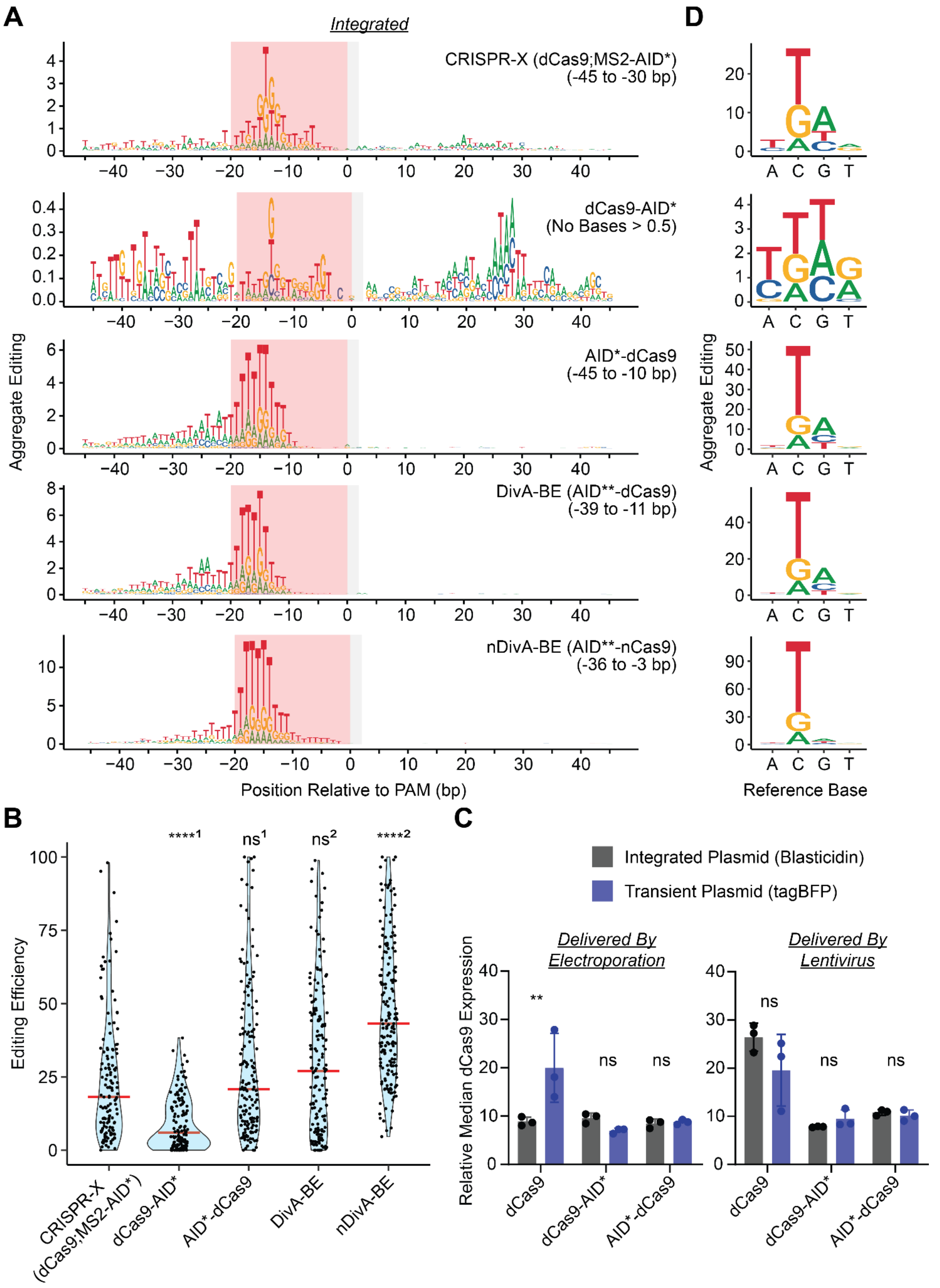

A
Integrated
CRISPR-X (dCas9;MS2-AID*)
(-45 to -30 bp)
dCas9-AID*
(No Bases > 0.5)
AID*-dCas9
(-45 to -10 bp)
DivA-BE (AID**-dCas9)
(-39 to -11 bp)
nDivA-BE (AID**-nCas9)
(-36 to -3 bp)
Aggregate Editing
Position Relative to PAM (bp)
D
Reference Base
B
Editing Efficiency
CRISPR-X (dCas9;MS2-AID*)
dCas9-AID*
AID*-dCas9
DivA-BE
nDivA-BE
C
Integrated Plasmid (Blasticidin)
Transient Plasmid (tagBFP)
Delivered By Electroporation
Delivered By Lentivirus
Relative Median dCas9 Expression
dCas9
dCas9-AID*
AID*-dCas9

**Supplementary Figure S3. Editing efficiency increase for dCas9-AID* was dependent on the delivery method**

(A) Aggregate editing logos of base substitutions produced by Integrated CRISPR-X (dCas9;MS2-AID*), dCas9-AID*, AID*-dCas9, DivA-BE (AID**-dCas9), nDivA-BE (AID**-nCas9) base editors at each base position relative to the PAM. The bases shown indicate the resulting base substitution. The red-shaded boxes indicate the position of the protospacer sequence for sgRNAs. The gray-shaded boxes indicate the position of the NGG PAM.

(B) Violin plots for editing efficiency of Integrated CRISPR-X, dCas9-AID*, AID*-dCas9, DivA-BE, nDivA-BE base editors. The red bar indicates the median editing efficiency.

(C) Relative dCas9 expression levels for Integrated and Transient dCas9, dCas9-AID*, and AID*-dCas9 expression vectors delivered by electroporation or lentivirus to wild-type K562 cells (n=3 biological replicates, one-way ANOVA test with Dunnett's multiple comparisons test; error bars represent standard deviation; ** - $p \leq 0.01$).

(D) Aggregate editing plot of editing spectrum of base substitutions generated by Integrated CRISPR-X, dCas9-AID*, AID*-dCas9, DivA-BE, nDivA-BE base editors. (Kruskal-Wallis test; [1] indicate comparisons to CRISPR-X and [2] indicate comparisons to AID*-dCas9; **** - $p < 0.0001$, ns - not significant)

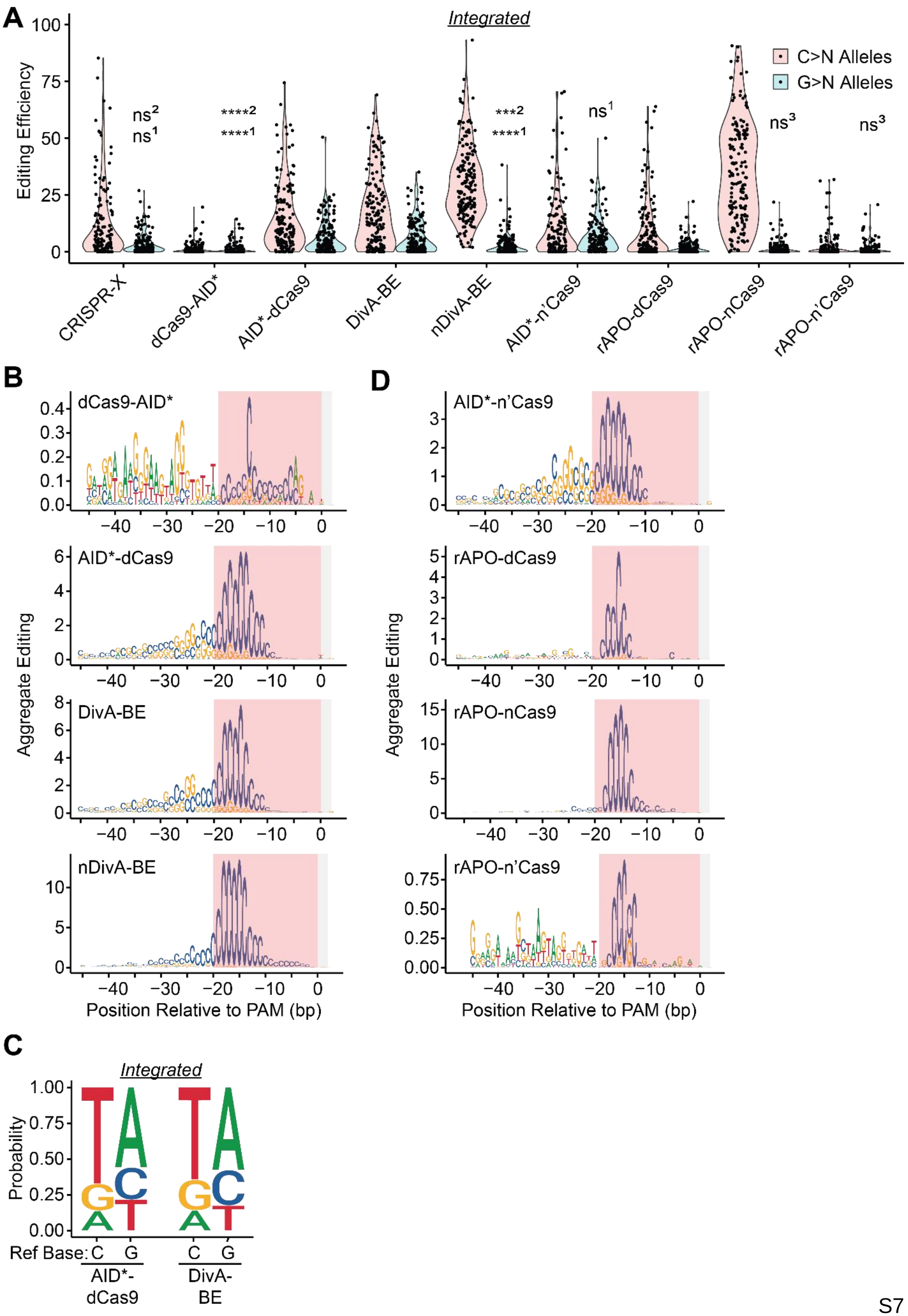

A
Integrated
Editing Efficiency
C>N Alleles
G>N Alleles
ns2
ns1
****2
****1
***2
****1
ns1
ns3
ns3
CRISPR-X
dCas9-AID*
AID*-dCas9
DivA-BE
nDivA-BE
AID*-n'Cas9
rAPO-dCas9
rAPO-nCas9
rAPO-n'Cas9
B
dCas9-AID*
AID*-dCas9
DivA-BE
nDivA-BE
Aggregate Editing
Position Relative to PAM (bp)
D
AID*-n'Cas9
rAPO-dCas9
rAPO-nCas9
rAPO-n'Cas9
Aggregate Editing
Position Relative to PAM (bp)
C
Integrated
Probability
Ref Base: C G C G
AID*-dCas9
DivA-BE

**Supplementary Figure S4. Nicking the non-target strand increased relative G>N editing for AID based editors**

(A) Violin plots for editing efficiency of Integrated CRISPR-X, dCas9-AID*, AID*-dCas9, DivA-BE, nDivA-BE, AID*-n'Cas9, rAPO-dCas9, rAPO-nCas9, and rAPO-n'Cas9 base editors. Red violin plots represent the editing of alleles containing C>N mutations, and blue plots indicate efficiency for G>N mutations. (Kruskal-Wallis test; [1] indicate comparisons to G>N editing for AID*-dCas9, [2] indicate comparisons to DivA-BE and [3] indicate comparisons to rAPO-dCas9; *** - $p \leq 0.001$, **** - $p < 0.0001$, ns - not significant)

(B) Aggregate editing logos of base substitutions produced by Transient dCas9-AID*, AID*-dCas9, DivA-BE, and nDivA-BE base editors at each base position relative to the PAM. The bases shown indicate the reference base that was altered. The red-shaded boxes indicate the position of the protospacer sequence for sgRNAs. The gray-shaded boxes indicate the position of the NGG PAM.

(C) Normalized editing spectra for C>N and G>N editing in AID*-dCas9 and DivA-BE samples for Integrated delivery.

(D) Aggregate editing logos of base substitutions produced by Integrated AID*-n'Cas9, rAPO-dCas9, rAPO-nCas9, and rAPO-n'Cas9 base editors at each base position relative to the PAM. The bases shown indicate the reference base that was altered. The red-shaded boxes indicate the position of the protospacer sequence for sgRNAs. The gray-shaded boxes indicate the position of the NGG PAM.

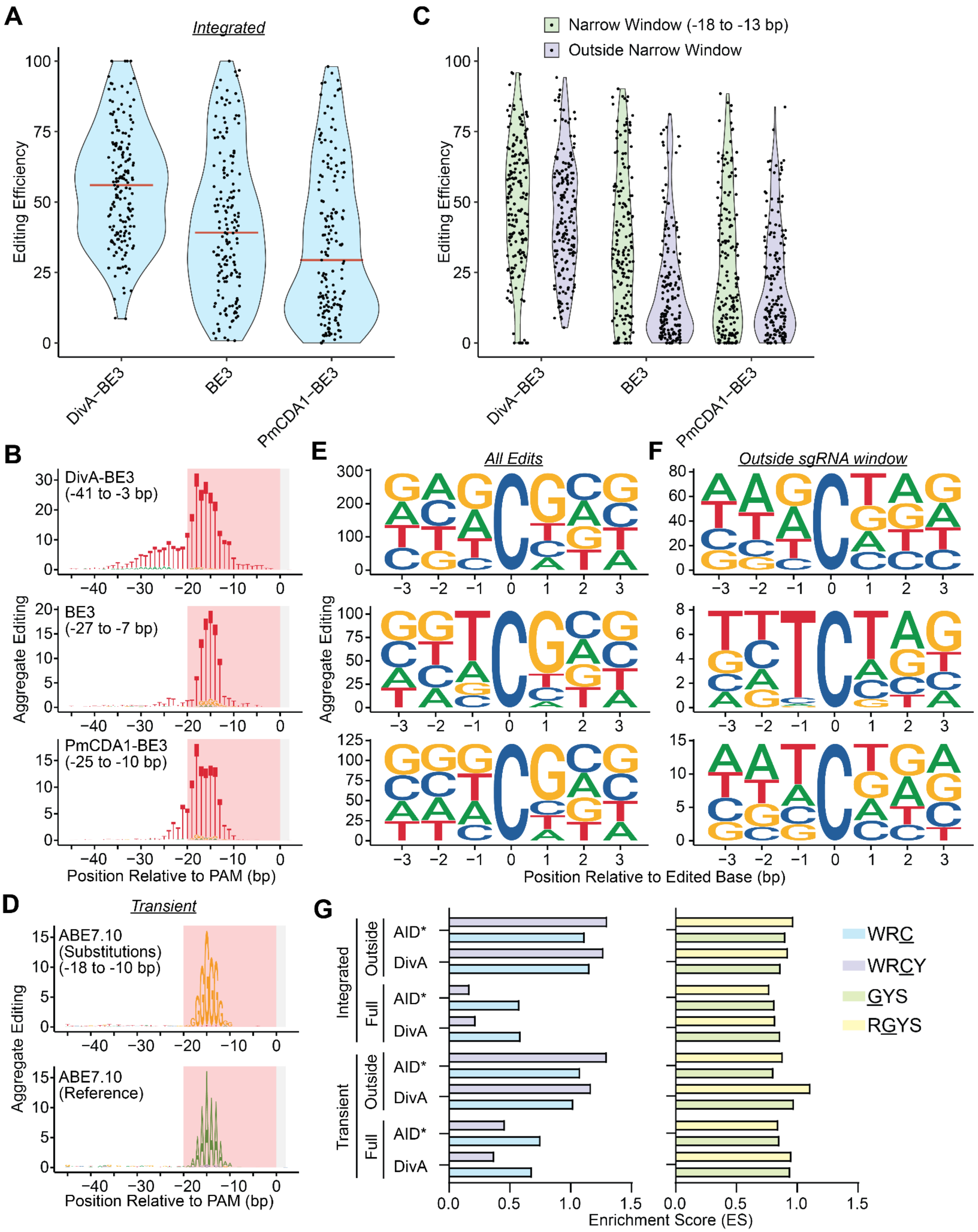


**Supplementary Figure S5. Cytidine and guanine motif editing preferences were enriched outside of the sgRNA targeting window**

(A) Violin plots for editing efficiency of Integrated DivA-BE3, BE3, and PmCDA1-BE3 base editors. The red bar indicates the median editing efficiency.

(B) Aggregate editing logos of base substitutions produced by Integrated DivA-BE3, BE3, and PmCDA1-BE3 base editors at each base position relative to the PAM. The bases shown indicate the resulting base substitution. The red-shaded boxes indicate the position of the protospacer sequence for sgRNAs. The gray-shaded boxes indicate the position of the NGG PAM.

(C) Violin plots for editing efficiency of Integrated DivA-BE3, BE3, and PmCDA1-BE3 base editors. Green violin plots represent the efficiency of alleles containing edits in the narrow window (-18 to -13 bp), and the purple plots indicate the efficiency of alleles containing edits outside the window.

(D) Aggregate editing logos of base substitutions produced by the Transient ABE7.10 base editor at each base position relative to the PAM. The top panel shows the resulting base from the substitution, and the bottom panel indicates the reference base that was edited. The red-shaded boxes indicate the position of the protospacer sequence for sgRNAs. The gray-shaded boxes indicate the position of the NGG PAM.

(E) Position-weight matrix for edited cytidines outside the -20 to 0 bp window for Transient DivA-BE3, BE3, and PmCDA1-BE3.

(F) Position-weight matrix for guanines and cytidines edited by Transient/Integrated DivA-BE3, BE3, and PmCDA1-BE3.

(G) Enrichment scores for motifs of edited cytidines or guanines. Scores are calculated for AID*-dCas9 and DivA-BE Transient and Integrated editors for indicated motifs considering edits across the editing window (Full) or those outside of -20 to 0 bp window (Outside).

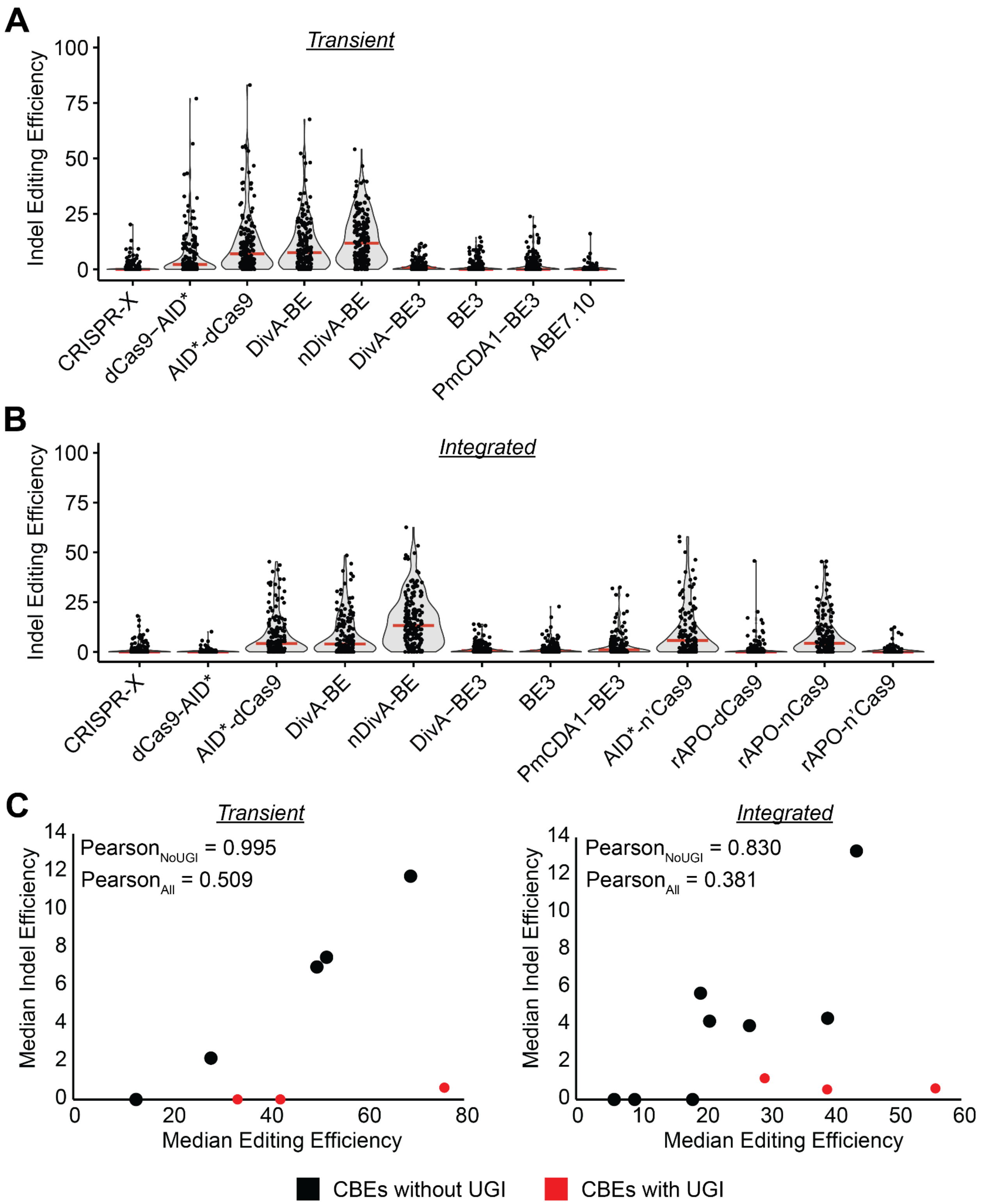


**Supplementary Figure S6. Base editors with a UGI exhibited a reduced indel efficiency relative to overall editing efficiency**

(A) Violin plots for indel editing efficiency of Transient single base editors tested in this study. The red bar indicates the median editing efficiency.

(B) Violin plots for indel editing efficiency of Integrated base editors tested in this study. The red bar indicates the median editing efficiency.

(C) Scatterplot of median indel and median editing efficiency for Integrated and Transient editors. Red circles indicate points for cytidine base editors (CBEs) that include a uracil glycosylase inhibitor (UGI). Pearson correlation between efficiencies for all editors ($Pearson_{All}$) and editors without a UGI ($Pearson_{NoUGI}$) is indicated on the graph.

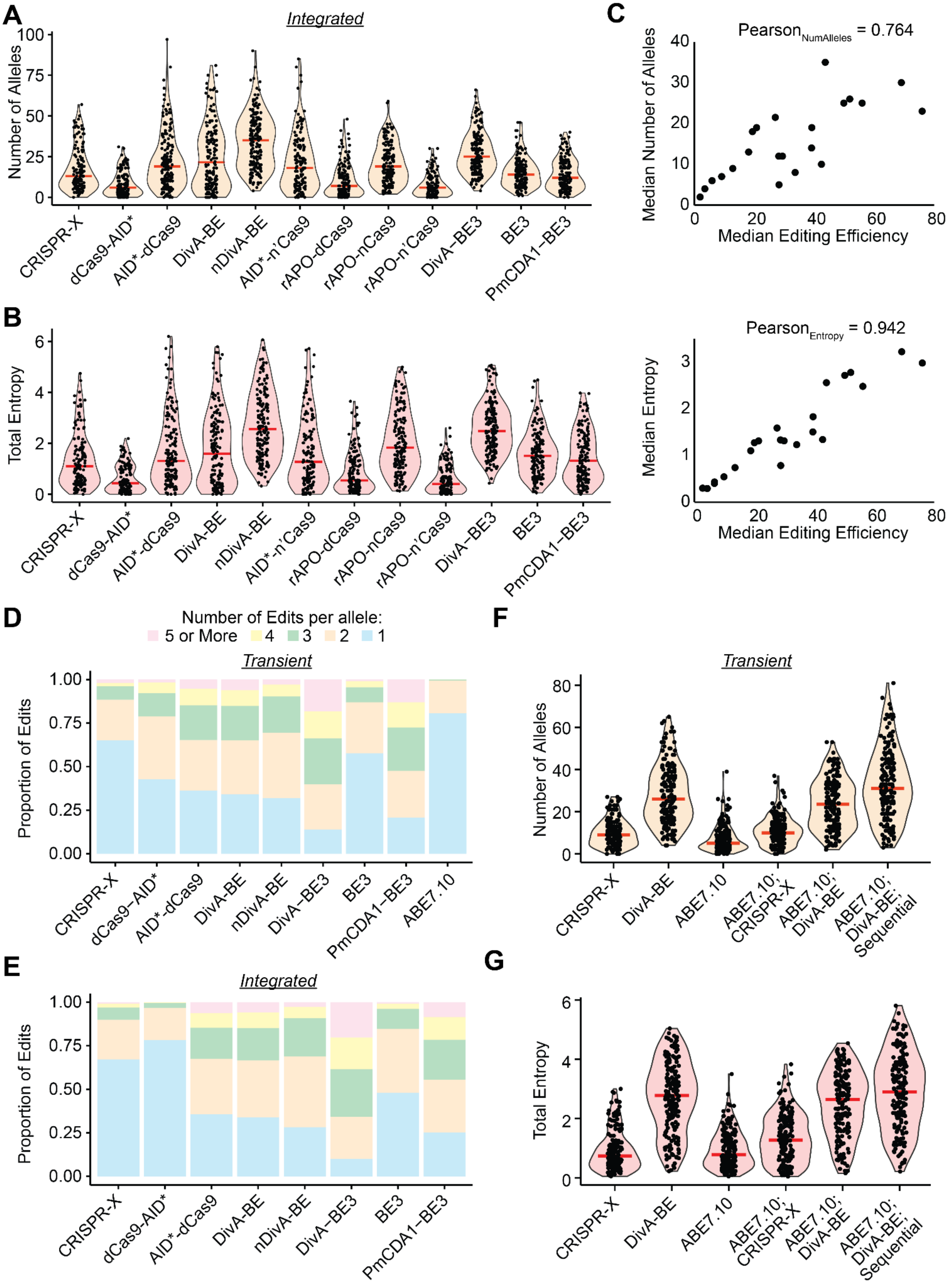
A
Integrated
Number of Alleles
CRISPR-X
dCas9-AID*
AID*-dCas9
DivA-BE
nDivA-BE
AID*-n'Cas9
rAPO-dCas9
rAPO-nCas9
rAPO-n'Cas9
DivA–BE3
BE3
PmCDA1–BE3
B
Total Entropy
C
Pearson$_{NumAlleles}$ = 0.764
Median Number of Alleles
Median Editing Efficiency
Pearson$_{Entropy}$ = 0.942
Median Entropy
D
Number of Edits per allele:
5 or More
4
3
2
1
Transient
Proportion of Edits
ABE7.10
E
Integrated
F
Transient
ABE7.10; CRISPR-X
ABE7.10; DivA-BE
ABE7.10; DivA-BE; Sequential
G

**Supplementary Figure S7. High-efficiency diversifying cytidine base editors produced alleles with multiple mutations**

(A) Violin plots for the number of alleles generated by Integrated base editors. Red bars indicate the median number of alleles generated.
(B) Violin plots for the total entropy generated by Integrated base editors. Red bars indicate the median entropy generated.
(C) Scatterplots for correlation between median editing efficiency and median number of alleles generated (top) and median entropy (bottom) for Transient and Integrated editors. Pearson correlation between values is indicated on the graph.
(D) Stacked bar plot of the proportion of edits with the number of mutations per allele for Transient base editors
(E) Stacked bar plot of the proportion of edits with the number of mutations per allele for Integrated base editors
(F) Violin plots for the number of alleles generated by a Transient combination of adenine and cytidine base editors: ABE7.10;CRISPR-X, ABE7.10;DivA-BE, and ABE7.10;DivA-BE;Sequential. Red bars indicate the median number of alleles generated.
(G) Violin plots for the entropy generated by a Transient combination of adenine and cytidine base editors: ABE7.10;CRISPR-X, ABE7.10;DivA-BE, and ABE7.10;DivA-BE;Sequential. Red bars indicate the median number of alleles generated.

**Supplementary Text S1**

**Supplementary Methods**

**Comparison of read filtering pipelines**

We compared four filtering pipelines by analyzing the Transient Parent, dCas9;MS2-AID* (CRISPR-X), dCas9;MS2-AIDDead, and AID*-dCas9 samples for both replicates. For each sample, fastq files after extraction and truncation were aligned as described in the main methods text using bowtie (sgRNA) and bowtie2 (target site) aligners. The 'Aligned' filtering method removed target site reads that failed to align. The 'Matched' filtering method removed reads where the aligned target site and sgRNA were improperly paired. This Matched filter was used for the additional two filtering methods. For the last two methods, we identified sgRNA-target site-UMI triplets in the Parent sample that passed a specified filter. For 'Matched-UMI-Indel' method, sgRNA-target site-UMI triplets with ≤ 90% of their reads not containing insertions/deletions were removed. For 'Matched-UMI-Perfect' method, sgRNA-target site-UMI triplets with ≤ 66% of their reads having perfect alignment were removed. The list of acceptable triplets for the 'Matched-UMI-Indel' and 'Matched-UMI-Perfect' were generated by the 'Define_Whitelist.py' script.

We filtered the fastq files from the samples using 'UMI-Filter_ProcComp.py'. This script generates four fastq files for each sample using the four filtering methods: Aligned, Matched, Matched-UMI-Indel, and Matched-UMI-Perfect. For the Matched-UMI methods, only reads that matched an acceptable sgRNA-target site-UMI triplet were retained in the processed fastq file. We processed the resulting fastq files using CRISPResso2 with the settings described in the methods in the main text. The outputs of all CRISPResso2 analyses were combined into a single table using 'CreateAlleleTables_General.py' followed by normalization of the tables using 'NormalizeTables_Initial_Cutoff.py'. This script removed alleles with only one read, and the frequency of each allele was recalculated.

For each filtering method, we calculated the normalized frequency of each allele detected in the CRISPR-X, dCas9;MS2-AIDDead, and AID*-dCas9 samples using 'MakeNormTables_Initial_Cutoff.r'. The normalization of the allele frequencies to the Parent sample and calculations of editing efficiency for each target site were performed as described in the methods of the main text.

**Calculating the correlation between indel and substitution frequencies**

The allele table generated by 'NormalizeTables_Initial_Cutoff.py' was filtered to analyze the allele frequencies generated by the Aligned filtering method for the Parent and dCas9;MS2-AIDDead Rep1 and Rep 2 samples. CRISPResso2 analysis outputs the number of bases that have substitutions (n_mutated), the number of inserted bases (n_inserted), and the number of deleted bases (n_deleted). These values were used to identify alleles that contain indels only (n_mutated = 0) and substitutions only (n_inserted and n_deleted = 0). Pairwise Pearson correlation coefficients between the allele frequencies in all four samples were calculated.

### Supplementary Text S2

### Supplementary Results

#### Sequencing sgRNA and filtering with UMI improved the detection of base editing

We compared four pipelines for filtering reads to detect base editing in our assay. For this comparison, we used four samples from the Transient set: Parent, AID*-dCas9, dCas9;MS2-AID* (CRISPR-X), and dCas9;MS2-AIDDead. In the first method (Aligned), we aligned the target site read without considering the UMI or sgRNA sequencing read. As expected, we detected reads containing "mutations" in the Parent sample, as mutations can be introduced at multiple steps, such as oligonucleotide synthesis, molecular cloning, lentiviral production, or sequencing library preparation. If the mutations were systematically introduced, the mutations should be detected at similar frequencies in non-Parent samples within the same replicate and could be normalized computationally. To analyze whether the errors were systematically being introduced, we calculated the correlation between allele frequencies in the Parent and dCas9;MS2-AIDDead samples across replicates (**Figure S1C**), neither of which should be edited. We analyzed alleles that contained indels only or substitutions only, as each type of mutation may be introduced at different steps in the sample processing pipeline. We observed a high correlation between Parent and dCas9;MS2-AIDDead samples within each replicate for indels (Pearson = 0.86 and 0.75 for replicates 1 (Rep1) and 2 (Rep2), respectively) and base substitutions (Pearson = 0.88 and 0.79 for Rep1 and Rep2). We observed a low correlation (Pearson = 0.25-0.34) for indels between replicates, while substitutions showed a more modest correlation (Pearson = 0.39-0.51).

Given the high correlation of indels and substitutions within replicates, we used the Parent sample to normalize the editing detected in samples that were exposed to genome editors. After this normalization, we calculated editing efficiency for all target sites in the AID*-dCas9, CRISPR-X, and dCas9;MS2-AIDDead Transient samples (**Figure S1D**). We expected no editing for the dCas9;MS2-AIDDead samples, while we expected AID*-dCas9 to have a higher editing efficiency than CRISPR-X based on tests with individual sgRNAs (data not shown). The negative control sgRNA-target site pairs were analyzed separately, as these sites should not be edited for all samples, including those with active AID*. To our surprise, we detected a slight increase in median editing efficiency for the two AID* samples (10.63% and 10.62%) compared to dCas9;MS2-AIDDead (7.48%). These findings suggested that we included many alleles that were not efficiently edited.

An explanation for the low editing efficiency could be recombination between sgRNAs and target sites during plasmid library construction and lentiviral transduction. These recombination events would result in mismatched sgRNA-target site pairs being delivered to a cell. This cell would be unable to edit since the sgRNA does not direct Cas9 to the target site. Removing reads with improperly paired sgRNAs and target sites should increase the measured editing efficiency, although it would also remove target site reads that were correctly matched in the cell when editing was induced but underwent recombination during PCR amplification when preparing the sequencing library preparation (1). We repeated the editing efficiency analysis only considering sequencing reads where the sgRNA and target site matched ('Matched' method) to remove improperly paired reads. This additional filtering increased median editing efficiency for the AID*-dCas9 (45.29%) and CRISPR-X (18.33%), consistent with our previous observation that AID*-dCas9 was a more efficient editor (**Figure S1D**). dCas9;MS2-AIDDead (8.79%) exhibited an increase in median editing compared to the Aligned method, but the increase was less substantial than the other editors. Based on this improved detection of editing, we included this Matched filter in the further downstream methods.

The Matched filtering method detected editing for negative control sgRNA-target site pairs for all three editors analyzed (6-12%) (**Figure S1D**). We suspected these background mutations could be derived from target sites harboring mutations before editing was induced. Therefore, we wanted to limit our analysis to acceptable target sites that did not contain mutations before editing was induced. We produced a list of acceptable sgRNA-target site-UMI triplets in the Parent sample, to which we would restrict our downstream analysis. Similar methods have been used to reduce background and increase detection sensitivity in assays profiling the activity of Cas9 nucleases (2). These previous applications of similar filtering methods found that errors in oligonucleotide synthesis frequently result in indels. Therefore, we applied a filter that removed these indels (Matched-UMI-Indel), which only considered sgRNA-target site-UMI triplets where > 90% of the reads did not contain an indel. An additional filtering method required that > 66% of the reads for a triplet be a perfect match (Matched-UMI-Perfect). The Matched-UMI-Indel filter reduced the median editing efficiency induced in negative control sgRNA-target site pairs for AID*-dCas9 (0%) and CRISPR-X (2.69%), respectively. The Matched-UMI-Perfect filtering method further reduced editing efficiency at negative control target sites, but over half of the On-Target sites had to be dropped from the analysis due to low read count. Therefore, we proceeded with the Match-UMI-Indel filtering for all subsequent analyses (workflow is in **Figure S1E**).
